# A functional trade-off between executive control and implicit statistical learning is dynamically gated by mind wandering

**DOI:** 10.1101/2025.08.05.668618

**Authors:** Teodóra Vékony, Bianka Brezóczki, Gábor Csifcsák, Dezső Németh, Péter Simor

## Abstract

Human cognition must balance goal-directed behavior with the need to learn from environmental regularities. Mind wandering (MW), a state of attentional decoupling from the task at hand, is paradoxically associated with both executive failures and enhanced implicit statistical learning, yet the direct relationship between these phenomena remains unclear. Here, we provide direct behavioral evidence for a functional trade-off between these competing demands. Using a task that concurrently measured response inhibition, statistical learning, and self-reported task focus, we show that MW is associated with impaired inhibitory control but enhanced learning of probabilistic sequences. Critically, we reveal that these effects are related: the magnitude of the learning enhancement during MW depends on the efficacy of response inhibition. These findings demonstrate that transient lapses in top-down executive control are associated with the enhanced implicit extraction of environmental statistics, supporting neurocompetition models, and framing MW as a cognitive state that may be evolutionarily preserved to promote the unsupervised acquisition of predictive models.

## Introduction

Mind wandering (MW) occurs when attention shifts away from the task at hand, and our thoughts drift towards internal reflections about the past or future. This state is typically linked to reduced cognitive performance in areas such as sustained attention (Seli et al., 2015), executive control (Andrillon et al., 2021; McVay & Kane, 2012), reading comprehension (McVay & Kane, 2012), model-based decision-making (Liu et al., 2023), and fluid intelligence (Mrazek, Smallwood, & Schooler, 2012; Mrazek, Smallwood, Franklin, et al., 2012). Behaviorally, these lapses are characterized by a complex signature of altered performance. While MW is frequently associated with lower accuracy and quicker, more impulsive responses indicative of executive failures (Mooneyham & Schooler, 2013), it is equally linked to unusually sluggish responses and increased overall reaction time variability (Bastian & Sackur, 2013; Boayue et al., 2021). Neurocognitively, this performance decrement is driven not only by a breakdown in top-down cognitive control but also by the phenomenon of perceptual coupling (Schooler et al., 2011), during which the brain attenuates the sensory processing of external environmental stimuli to shield internal trains of thought. Consequently, MW has been consistently associated with the attenuation of sensory processing, as reflected by behavioral measures and neural markers (Arnau et al., 2020; Dong et al., 2021; Kam et al., 2011, 2013). Recent research suggests that these lapses may stem from the emergence of localized, sleep-like slow waves within the awake brain. Specifically, frontal slow waves have been shown to predict MW and impulsive behaviors, providing a physiological explanation for the transient intrusion of task-unrelated thoughts (Andrillon et al., 2021).

Despite the drawbacks and the withdrawal of sensory and executive resources, humans spend a significant portion of their time in a MW state (Bastian & Sackur, 2013; Cheyne et al., 2009; Mooneyham & Schooler, 2013). This suggests that MW might serve some adaptive purpose. Recent studies show that MW can provide cognitive benefits, particularly under low attentional demands and when the goals of learning are opaque or fully implicit (Amer et al., 2016; Thompson-Schill et al., 2009), such as in the case of statistical learning (Simor et al., 2025; Vékony et al., 2025). However, the exact mechanism behind this benefit remains unclear. Here, we test the hypothesis that the transient state of executive failure during MW is associated with the implicit extraction of statistical regularities from the environment. To arbitrate this, we employ a paradigm designed to simultaneously capture the dynamics of MW, response inhibition, and statistical learning, thereby offering a unified account of the cognitive trade-offs between endogenous thought and statistical learning.

Research has shown that MW significantly impacts executive functioning, including response inhibition, which governs quick behavioral action restriction or action cancellation (Kam & Handy, 2014; Maillet et al., 2020). This effect often manifests as an inability to stop a prepotent response, indicating that top-down control has temporarily broken down. Despite its deleterious impact on executive functions, MW has been repeatedly linked to our ability to discern complex statistical patterns in our environment. A paradox lies at the heart of human cognition: attenuated executive control is associated with both the rise of MW (Randall et al., 2014) and enhanced statistical learning (Simor et al., 2025; Vékony et al., 2025). This raises the possibility of a shared underlying mechanism. The two-sided nature of MW (marked by a negative association with executive control and a positive one with statistical learning) aligns with both theoretical considerations and empirical findings supporting the neurocompetition model (Ambrus et al., 2020; Nemeth et al., 2013; Smalle et al., 2017; Virag et al., 2015). This model posits that effortful, controlled processes and implicit, associative, and habitual learning compete for shared neurocognitive resources. While MW is often characterized by a shift of executive resources from external goals to internal thoughts (Smallwood et al., 2012a; Smallwood & Schooler, 2006), neurocompetition models suggest that this redirection of top-down control may paradoxically benefit certain forms of learning. Specifically, the withdrawal of effortful, on-task attention reduces the inhibitory influence that goal-directed processes typically exert over the implicit system. Accordingly, MW may shield executive control from effortful, task-specific processing, thereby facilitating implicit learning (Verschooren & Egner, 2023). Although the neurocompetition model has inspired a range of studies, empirical evidence for competition between cognitive processes remains largely indirect: executive functions, statistical learning, and their associations with MW are typically examined in separate tasks. The critical question of whether the attentional deficits associated with MW are accompanied by corresponding gains in implicit statistical learning can only be addressed by a single, integrated task to enable assessing the interplay between these cognitive phenomena on an intra-individual level. Here, we provide a direct empirical exploration of this complex interplay, aiming to elucidate the dynamics between these fundamental cognitive processes.

Healthy participants completed the Cognitive Trade-off Task (CTT), a combined inhibition and statistical learning task with thought probes inserted between task blocks. Consistent with prior work, our results confirm that periods of MW are characterized by compromised response inhibition and enhanced extraction of statistical regularities. Our principal novel finding, however, is the link between these phenomena. We show that the relationship between MW and statistical learning depends on response inhibition. This suggests a functional trade-off, where the transient lapse in top-down executive control during MW facilitates the implicit statistical learning of environmental patterns.

## Methods

### Participants

Three hundred and seventeen university students (249 females) were enrolled in the study. The mean age of the participants was 22.02 ± 4.12 SD years. After exclusions (see “*Quality control of the data*” section), two hundred and forty participants remained in the final sample. Participants received course credit for taking part in the study. Participants provided informed consent before enrolling. The experiment was conducted in accordance with the Declaration of Helsinki. The present study was not pre-registered. To ensure robust statistical power, our strategy relied on recruiting a large initial cohort of participants. The appropriateness of the final sample size was formally verified post-data collection through a simulation-based power analysis, detailed in the Statistical analysis section.

### Quality control of the data

Data integrity was ensured through a quality control examination (Rodd, 2024) based on predefined criteria (similar to Vékony et al., 2025). Participants were excluded if they reported a history of neurological or psychiatric disease (*n* = 43), use of nervous system-affecting medication (*n* = 8), alcohol consumption within three hours before the experiment (*n* = 19), or prior completion of the task (*n* = 10). Additionally, those who showed extreme non-compliance were disqualified on the basis that (1) task accuracy was less than 80% (*n* = 11), (2) task completion duration was greater than 50 minutes (*n* = 7), or (3) task restart occurred at any moment (*n* = 3). Seventy-seven individuals were excluded, some of whom satisfied more than one exclusion criterion.

### The Cognitive Trade-off Task (CTT)

The CCT is based on the Alternating Serial Reaction Time (ASRT) task (Howard & Howard, 1997), incorporating certain ‘No-Go’ trials to which participants must refrain from responding. The experimental task was programmed in JavaScript using the jsPsych library v.6.1.0 (de Leeuw, 2015). Participants responded to a visual stimulus—a dog’s or a cat’s head presented in one of four horizontal screen locations—by pressing the corresponding key (‘S’, ‘F’, ‘J’, or ‘L’ from left to right) (Figure 1A). The visual stimulus signaled either a ‘Go’ trial, requiring a fast key press response corresponding to the location of the stimulus, or a ‘No-Go’ trial, instructing participants to withhold any key press. On ‘Go’ trials, a correct key press led to the immediate disappearance of the stimulus, followed by a 120 ms interstimulus interval before the subsequent stimulus. In contrast, an incorrect response on a ‘Go’ trial meant the stimulus persisted until the correct key was pressed. For ‘No-Go’ trials, the stimulus was displayed for a fixed duration of 1000 ms, irrespective of any input from the participant.

**Figure 1.**
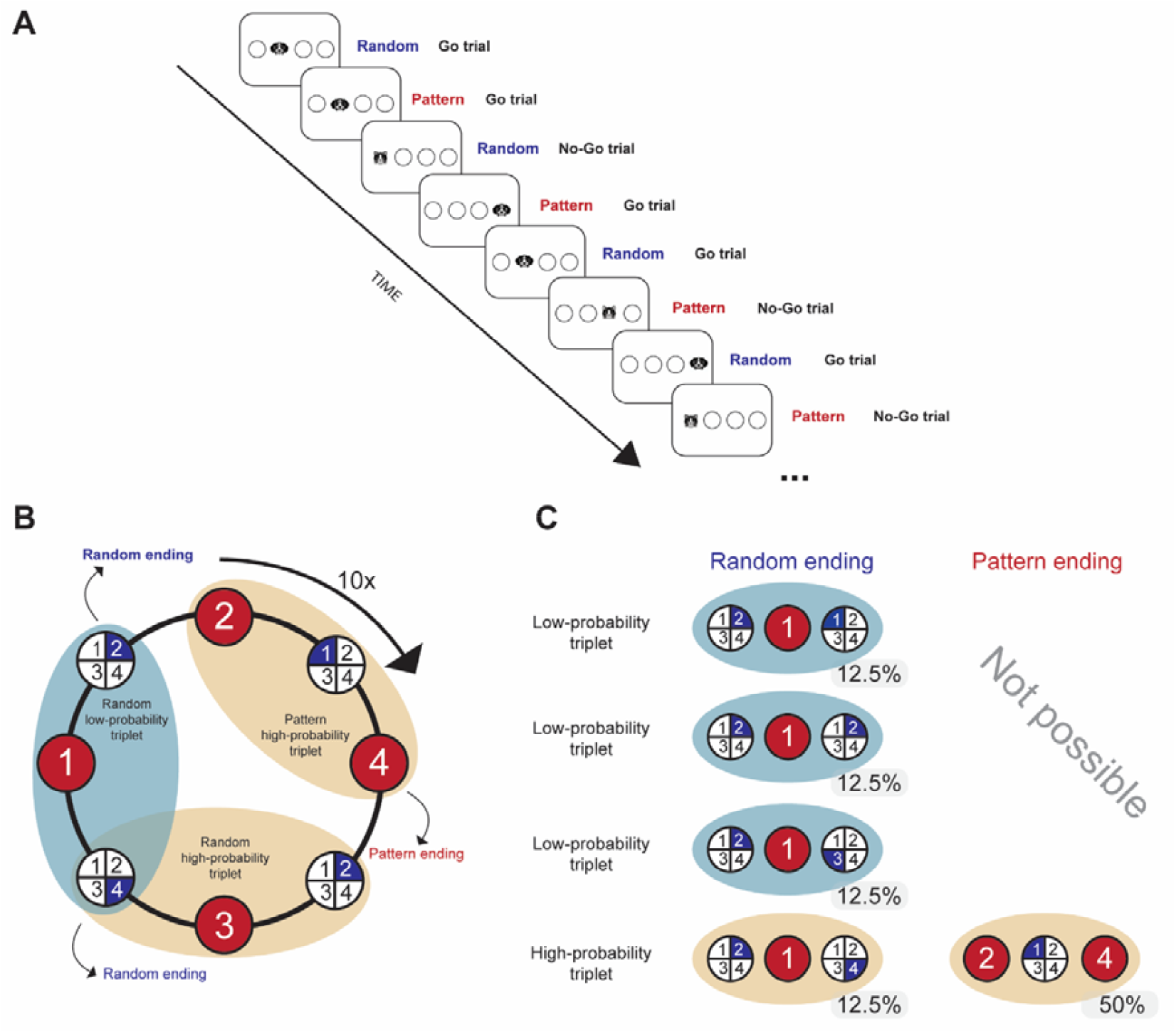
The Cognitive Trade-off Task. **(A)** Participants’ primary objective was to press the key that matched the on-screen location of a target stimulus (‘Go’ trial). Occasionally, a ‘No-Go’ stimulus appeared instead of the ‘Go’ stimulus, and participants had to withhold their response to these trials (‘No-Go’ trial). A key feature of this task was the underlying structure of the stimulus presentation: every second trial was part of an 8-element probabilistic sequence. The ‘No-Go’ stimuli could be either a pattern or a random element. Participants completed 80 trials in a block, of which 10 were ‘No-Go’ stimuli. **(B)** Participants encountered sequences of three consecutive trials, referred to as “triplets”. Elements could be either a pattern (red background) or random elements (blue background). Pattern elements consistently appeared in the same position throughout the task. In contrast, random elements were randomly chosen from four possible positions for each occurrence. The design of the task led to a probabilistic sequence structure, resulting in certain triplets appearing more frequently (high-probability triplets) than others (low-probability triplets). **(C)** The formation of high-probability triplets primarily involved two main configurations. In 50% of the trials, the triplet consisted of two pattern trials and one random trial positioned in the center. Another configuration, occurring in 12.5% of trials, involved two random trials and one pattern trial at the center, also forming a high-probability triplet. Combined, these configurations meant that 62.5% of all trials concluded with a high-probability triplet, while the remaining 37.5% were the final elements of a low-probability triplet. Panel B and C was reprinted with permission from Figure 1 of Vékony et al. 2025, *iScience*, which was published under a CC BY-NC-ND [https://creativecommons.org/licenses/by-nc-nd/4.0/]. Further reproductions must adhere to the terms of this license.

The stimuli (including both ‘Go’ and ‘No-Go’ trials) followed a probabilistic eight-element sequence, alternating between predictable (pattern) and unpredictable (random) elements (e.g., 2 – r – 4 – r – 3 – r – 1 – r, where ‘r’ denotes a random location and numbers indicate the four positions from left to right) (Figure 1B). Each participant completed the task with one of 24 repeating eight-element sequences across 30 blocks, with each block containing ten repetitions. Short breaks and thought probes followed each block.

The CTT employed a probabilistic sequence structure where certain three-element sequences (triplets) occurred with higher probability (high-probability triplets) than others (low-probability triplets). A single presentation within the sequence, whether pattern or random, constituted a trial and, critically, could also be the final element of a high-or low-probability triplet (high-vs. low-probability trials). The analysis focused on whether a trial concluded a high-or low-probability triplet, irrespective of its pattern or random designation within the alternating sequence. For instance, in the sequence 2 – r – 4 – r – 3 – r – 1 – r, triplets like 2-X-4, 4-X-3, 3-X-1, and 1-X-2 (where X is the middle element) were high-probability, while triplets like 2-X-1 or 2-X-3 were low-probability (Figure 1B; note that Figure 1B illustrates three example triplets: a pattern-ending high-probability triplet [2(P)-1(R)-4(P)], a random-ending high-probability triplet [2(R)-3(P)-4(R)], and a random-ending low-probability triplet [4(R)-1(P)-2(R)]; however, every three consecutive elements formed either a high-or low-probability triplet. For example, within the sequence 2(P)-1(R)-4(P)-2(R)-3(P)-4(R)-1(P)-2(R), six triplets are formed starting with the first pattern element: 2(P)-1(R)-4(P), 1(R)-4(P)-2(R), 4(P)-2(R)-3(P), 2(R)-3(P)-4(R), 3(P)-4(R)-1(P) [all high-probability], and 4(R)-1(P)-2(R) [low-probability]. Subsequent references to triplet type will focus on trials serving as the final element of these high-or low-probability triplets).

Across the task, 64 distinct triplets were possible (16 high-probability, 48 low-probability). High-probability triplets comprised either two pattern trials with a central random trial (50% occurrence) or two random trials with a central pattern trial (12.5% occurrence) (Figure 1C). Overall, 62.5% of trials were the final element of a high-probability triplet (high-probability trials), and 37.5% were the final element of a low-probability triplet (low-probability trials).

Each block of the CTT consisted of 70 ‘Go’ trials and 10 ‘No-Go’ trials. We randomly assigned the position of the ‘No-Go’ trials in each block, ensuring an equal distribution between random and pattern trials. Additionally, we ensured that two ‘No-Go’ stimuli never appeared consecutively, and a ‘No-Go’ stimulus was never among the first two trials. ‘Go’ and ‘No-Go’ trials were distinguished by different visual stimuli: a dog’s head or a cat’s head. The mapping of dog and cat heads to the ‘Go’ trial condition was counterbalanced across all participants and balanced between those who reported a preference for dogs versus cats to control for potential preference-related biases.

### Thought probes

After each block of the CTT, we asked participants to report on their mental state. While our primary goal was to investigate if MW, defined as task-unrelated thought, was associated with response inhibition and implicit statistical learning, we also explored three specific aspects: MW episodes with reportable content in contrast to “mind blanking” (MB, off-task periods with no reportable mental content), spontaneous versus deliberate manifestations of MW, as well as MW with positive vs. negative emotional content.

To achieve this, participants answered four questions after each block, designed to differentiate between these aspects. The first question (Q1) asked, “*To what degree were you focusing on the task during the previous block?*” with responses ranging from 1 (Not at all) to 4 (Completely). A score of 1 indicated their thoughts were completely elsewhere (e.g., friends, weekend plans), while a 4 meant they were entirely focused on the task. Participants were instructed to use options 2 and 3 if they believed their response would fall between 1 and 4.

If the answer to Q1 was either 1 or 2 (indicating a MW state), the following second question (Q2) appeared: “*When you were not focusing on the task, were you thinking of something in particular or just thinking about nothing?*” (1 - I was thinking about nothing; 4 - I was thinking about something in particular). A score of 1 meant their mind had wandered, but they were not thinking of anything specific, whereas a 4 meant they were thinking about something concrete (e.g., a book, recent events, task difficulty, or hunger). The third question (Q3) asked, “*Were you deliberate about where you focused your attention, or did it happen spontaneously?*” (1 - I was completely spontaneous; 4 - I was completely deliberate). A score of 1 suggested their attentional focus was effortless, while a 4 indicated they consciously directed their attention. The fourth question (Q4) asked, “*Were your thoughts more positive or more negative?*” (1 - Completely positive; 4 - Completely negative). A score of 1 indicated the thoughts were completely positive, while a 4 indicated their thoughts were completely negative.

If the answer to Q1 was either 3 or 4 (indicating an off-task state), instead of the three questions about the features of MW, control questions were posed (“*Did you focus more on speed or accuracy in the previous block?*”, “*How difficult was it for you to concentrate on the task in the previous block?*” and “*How tiring did you find the task?*”).

We opted for these four questions instead of a single, complex thought probe to avoid overwhelming participants and to gain a more nuanced understanding of their mental states. Participants selected their response by pressing a key from 1 to 4 on their keyboard. Results related to Q2-Q4 can be found in the Supplementary Material in sections S3, S4, and S5.

### Procedure

The experiment was executed via the Gorilla platform for online experimentation (https://gorilla.sc). Participants first provided informed consent, followed by a detailed explanation on the interpretation of the MW probes and the evaluation of their mental attention in various contexts (identical as described in Vékony et al., 2025). Participants thereafter undertook a concise quiz to evaluate their understanding and received the assessment. Following the quiz, participants completed 30 blocks of the CTT. After each block, participants responded to the thought probes and received feedback on their average reaction time (RT) and accuracy. After the task, participants were asked if they noticed any patterns and to describe them if they did. None of the participants could accurately define the alternating pattern integrated into the task. Participants then provided demographic information (e.g., age, gender, education) as well as responded to questions about their consumption and conditions.

### Statistical learning and visuomotor performance scores

For the CTT, each ‘Go’ trial was categorized as the final element of either a high-or low-probability triplet based on the two preceding trials. To ensure robust analysis, trills (e.g., 1-2-1) and repetitions (e.g., 2-2-2) were removed to prevent the contamination of statistical learning scores by pre-existing motoric and perceptual response tendencies (Song et al., 2007). Specifically, these patterns often elicit faster or slower responses due to mechanisms independent of the underlying sequence structure, which can lead to an overestimation or distortion of the true learning effect (Vaquero et al., 2006). The first two trials of each block were also excluded as they could not form a complete triplet; and trials with RT falling outside ±3 times the individual median absolute deviations, or those below 100 ms or above 1000 ms, were similarly discarded. Following the removal of outliers, the RT data remained marginally right-skewed; thus, square-root transformation was applied to enhance normality. We defined two key types of learning: *statistical learning*, operationalized as the difference in RT between high-probability and low-probability trials (referred to as the statistical learning score), and *visuomotor performance*, which encompassed the overall RT on the task and how these changed over time, irrespective of item probability.

In line with standard operational definitions in behavioral paradigms, the RT difference between high-and low-probability trials was used as the primary metric of *statistical learning*. While we acknowledge the classical theoretical distinction between learning (acquisition) and performance (expression) (Vékony et al., 2020, 2022), the ASRT task involves continuous and dynamic probabilistic sequences where these processes are profoundly intertwined. Therefore, our metric of statistical learning is conceptualized as capturing both the ongoing, active extraction of environmental regularities and their concurrent behavioral expression.

### No-Go performance score

We calculated the percentage of correct ‘No-Go’ trials in each block to evaluate inhibitory control performance. A response was categorized as correct if no response key was pressed during the 1000-ms-long presentation of the ‘No-Go’ stimulus.

### MW scores

A dichotomous MW variable was created by categorizing the responses to thought probe Q1 into MW (off-task) (1–2) and on-task (3–4) periods. Contrasts between MW with identifiable content vs. MB were created by dichotomizing responses to Q2 (1–2 vs. 3–4). Similarly, we contrasted task performance between spontaneous vs. deliberate MW by categorizing responses to Q3 (1–2 vs. 3–4) and between positive vs. negative MW by categorizing responses to Q4 (1-2 vs. 3-4) for thought probes indicating MW for Q1.

### Statistical analysis

Statistical analysis was conducted in *R* (4.2.3). Simple regressions utilized the *lm* function from the *lme4* package, while linear mixed models (LMMs) were fitted using the *mixed* function (from *afex*) with sum-to-zero contrasts. For LMMs, we began with a maximal random-effects structure and simplified it to achieve convergence by removing correlations between random slopes or the slopes themselves (Meteyard & Davies, 2020). Numerical fixed factors were mean-centered. Fixed effects were evaluated using Type III tests.

Estimated marginal means and estimated marginal trends for post-hoc simple means and simple slopes analyses were calculated with the *emmeans* package, utilizing the default Tukey method to correct for multiple comparisons. Significant interactions involving continuous predictors were examined using estimated marginal trends to directly analyze continuous linear slopes. Specifically, when an interaction involved two continuous variables simultaneously (i.e., Block and No-Go performance), the continuous linear trend of the Block predictor was evaluated at standardized levels of the No-Go performance predictor, namely, the mean, as well as one standard deviation below (-1 SD) and above (+1 SD) the mean. All analyses used an alpha level of .05. Figures were generated with *ggplot2*, supplemented by *sjPlot, cowplot, ggplot*, and *afex* packages.

We conducted simple regressions to examine the evolution of MW during learning. The mean MW scores derived from raw 1-4 responses and the proportion of participants engaged in MW (MW vs. on-task periods) were our outcome variables, and Block (1-30) was a factor.

We performed a linear mixed model (LMM) to evaluate MW’s impact on response inhibition. The outcome variables were accuracy on ‘No-Go’ trials (in percentage), calculated per participant for each block. Fixed effects in this LMM included Block (1–30), MW (MW vs. on-task periods), and their interactions.

To understand the impact of MW on statistical learning and visuomotor performance, we also applied LMMs. The outcome variables were mean RT on ‘Go’ trials, calculated per participant for each block. To differentiate the assessment of statistical learning from response inhibition, the analysis of statistical learning on ‘Go’ trials primarily focused on RT (see analysis on accuracy in S2 section of the Supplementary Materials). This approach helped to avoid confounding measures, given that accuracy on ‘No-Go’ trials was specifically used to evaluate inhibitory control. First, we ran a model with fixed effects including Block (1–30), Triplet Type (last element of high-vs. low-probability triplets) and their interactions. Next, we ran another model also including MW (MW vs. on-task periods) and its interactions. We performed a more complex model by adding response inhibition performance as a continuous variable to the model (the percentage of correct ‘No-Go’ responses) along with their interactions. Model comparisons were performed using the *anova*() function and comparing the Akaike Information Criterion (AIC) values. Please note that Triplet Type main effects and interactions signal differences in *statistical learning*, while main effects and interactions without Triplet Type reflect differences in *visuomotor performance*. Full model tables can be found in the S1 section of the Supplementary Materials.

Finally, to confirm the adequacy of our sample size, we conducted a simulation-based power analysis for our main LMM using the *simr* package in R. Rather than calculating observed power, we established a Smallest Effect Size of Interest (SESOI) corresponding to a standardized small effect (Cohen’s d = 0.20), scaled to match the residual variance of our data. We specifically targeted the critical three-way interaction (Triplet Type × MW × No-Go Performance) for this simulation, as it represents the central theoretical hypothesis. Based on 1000 simulations, our design achieved 95.30% power (95% CI: 93.80%, 96.53%) to detect this minimal meaningful effect, confirming that the experiment was sufficiently powered.

## Results

### MW increases with task progression

A simple linear regression revealed a relationship between the progression of the task (Block 1–30) and participants’ mean MW scores. The model was statistically significant, *F*_(1, 7188)_ = 507.5, *p* < .001, explaining a significant portion of the variance in MW scores (adjusted R^2^ = .065). Specifically, task Block was a significant negative predictor of MW scores, *b* = −0.028, *t*_(7188)_ = −22.53, *p* < .001. This result indicates that as participants progressed through the task, their scores tended to decrease (indicating more MW) (Figure 2A).

**Figure 2.**
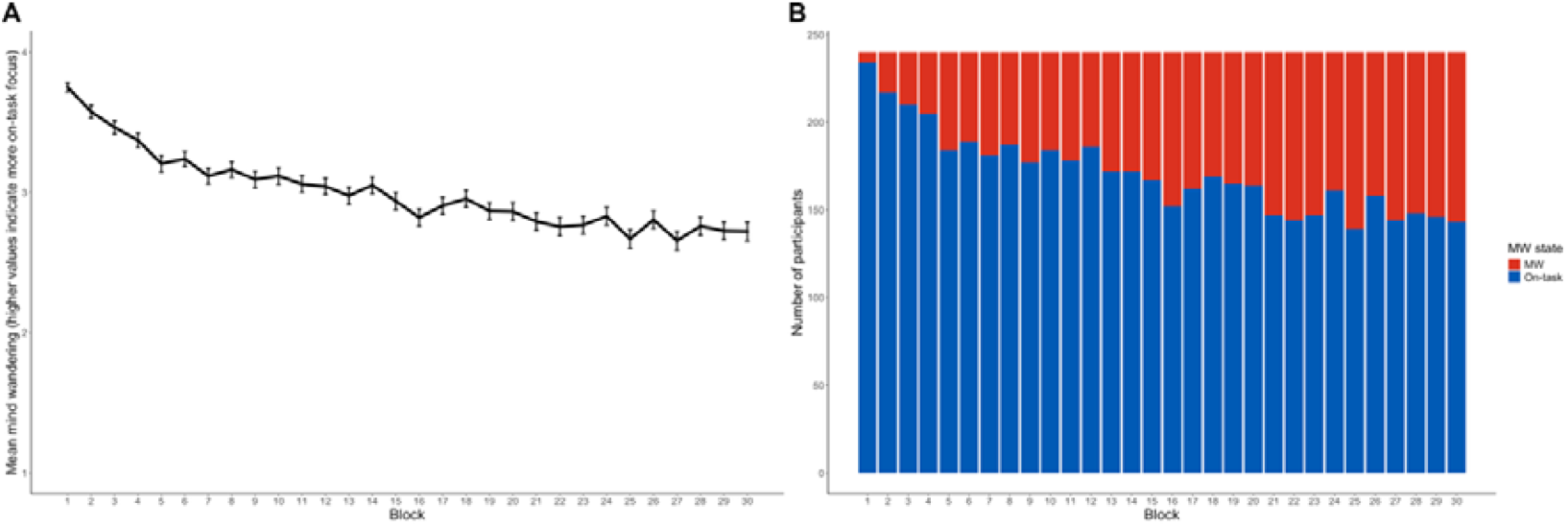
Change in MW throughout the CTT. (A) Mean MW score per block, as reported by participants. The x-axis represents block number, and the y-axis shows the average MW score (on a scale of 1–4, where lower scores indicate more MW). Error bars indicate SEM. (B) Number of participants engaged in MW per block. The x-axis indicates block number, while the y-axis reflects the number of participants. Stacked bars differentiate between participants who reported MW (red) and those who reported on-task focus (blue).

A simple linear regression was performed to examine the relationship between task Block (1– 30) and the proportion of participants engaged in MW (0–100%). The analysis revealed a statistically significant relationship, *F*_(1, 28)_ = 140.4, *p* < .001. The model accounted for a substantial amount of the variance in MW engagement, with an adjusted R^2^ of .83. The regression coefficient for Block (*b* = 0.010, *t*(28) = 11.849, *p* < .001) indicated a positive relationship: as the task progressed, the proportion of participants engaged in MW increased (Figure 2B).

### Response inhibition is impaired during periods of MW

An LMM was conducted to examine the effects of MW on response inhibition (defined as ‘No-Go’ accuracy), incorporating a by-participant random slope for Block (1-30), MW (MW vs. on-task), and their interaction to account for individual variability. The analysis revealed significant main effects for both MW (*F*_(1, 183.02)_ = 143.61 *p* < .001) and Block (*F*_(1, 212.98)_ = 152.19, *p* < .001), indicating that response inhibition significantly differed across levels of both factors. More specifically, response inhibition was worse during MW periods (*b* = - 0.046, *SE* = 0.004) and significantly decreased as the task progressed (*b* = -0.053, *SE* = 0.004) (Figure 3). However, the interaction between MW and Block was not statistically significant (*F*_(1, 155.68)_ = 0.140, *p* = .709), suggesting that the effect of MW on response inhibition did not change with the progress of the task.

**Figure 3.**
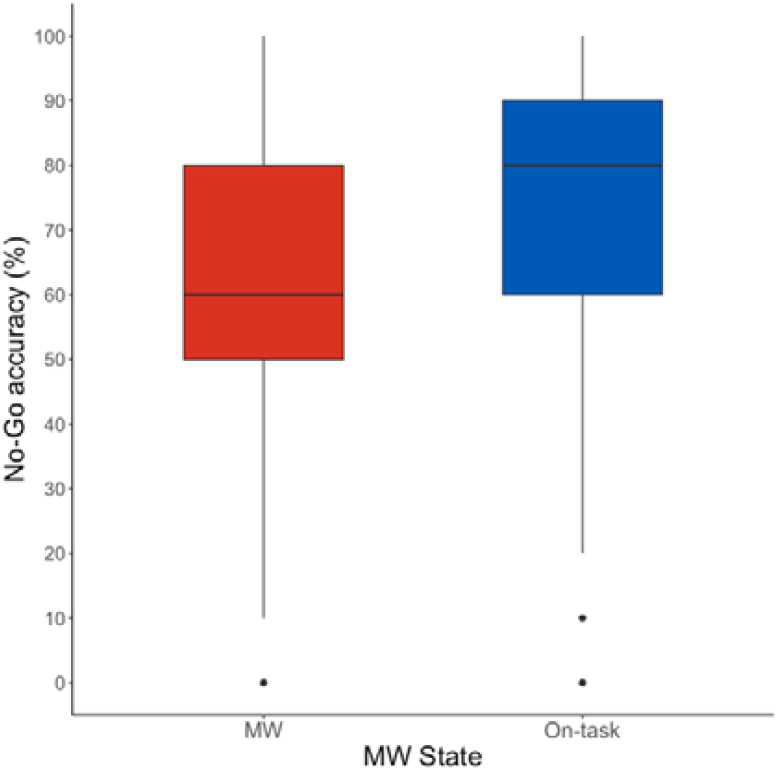
No-Go accuracy in the CTT. The x-axis represents the MW state reported by participants (MW or on-task), and the y-axis shows the mean accuracy on the ‘No-Go’ trials in a specific block. During periods of on-task states, participants performed significantly better on the ‘No-Go’ trials, i.e., they were more successful in inhibiting the keypresses. Whiskers denote 1.5 times the interquartile range, while outliers beyond this range are depicted as dots.

### Statistical learning is observed in the CTT

A LMM was conducted to evaluate whether statistical learning occurred in the CTT. Block (1-30) and Triplet Type (the last element of a high-vs. low-probability triplet) were fixed effects in the model. The dependent variable was RT on ‘Go’ trials.

The model converged with by-participant correlated random slopes for the Block and Triplet Type factors. The main effect of Block was significant (*F*_(1, 236.62)_ = 208.26, *p* < .001), suggesting faster RTs as the task progressed (*b* = -0.402, *SE* = 0.028).

The main effect of Triplet Type (*F*_(1, 227.88)_ = 110.44, *p* < .001) indicated overall faster RTs for the last element of high-probability triplets (*b* = -0.079, *SE* = 0.007), signaling statistical learning. The interaction between Block and Triplet Type was also significant (*F*_(1, 13626.04)_ = 17.36, *p* < .001). Post-hoc analysis of estimated marginal trends revealed that RT decreased at a significantly steeper rate across blocks for high-probability trials (*b* = -0.431, *SE* = 0.029) compared to low-probability trials (*b* = -0.373, *SE* = 0.029), signifying that statistical learning progressively improved as the task advanced.

### Statistical learning in the context of MW

As we observed that statistical learning did occur in the CTT, we examined how MW (MW vs. on-task) periods interacted with statistical learning. An LMM was conducted with MW (MW vs. on-task) as an additional fixed effect in the model. The dependent variable was RT on ‘Go’ trials.

The model converged with by-participant correlated random slopes for the Block and MW factors and their interaction. The AIC comparison of the models with (*AIC* = 37206) and without the MW factor (*AIC* = 37661) indicated better goodness of fit for the more complex model (χ2_(8)_ = 470.629, *p* < .001). Triplet Type and Block main effects, as well as their interaction, appeared similarly as in the simpler model (see Supplementary Material S1.3 for details).

A main effect of MW appeared (*F*_(1, 113.26)_ = 12.37, *p* < .001), indicating faster RTs during periods of MW (*b =* -0.084, *SE* = 0.024). The interaction between MW and Block was also significant (*F*_(1, 139.75)_ = 8.50, *p* = .004). Post-hoc analysis of estimated marginal trends revealed that the decrease in RT across blocks was significantly less steep during MW periods (*b* = -0.318, *SE* = 0.038) compared to on-task periods (*b* = -0.405, *SE* = 0.024), indicating that the effect of MW on general RT was stronger at the beginning of the task and progressively diminished.

Most importantly, the interaction between Triplet Type and MW was also significant (*F*_(1, 13247.73)_ = 5.75, *p* =.016), suggesting that the magnitude of statistical learning between MW and on-task periods differed. Analysis of post-hoc estimated marginal means revealed a larger RT difference between high- and low-probability trials during periods of MW (*b* = - 0.206, *SE* = 0.027) than during on-task periods (*b* = -0.130, *SE* = 0.016) (Figure 4). No additional interactions concerning the MW factor were significant (*p* > .155).

**Figure 4.**
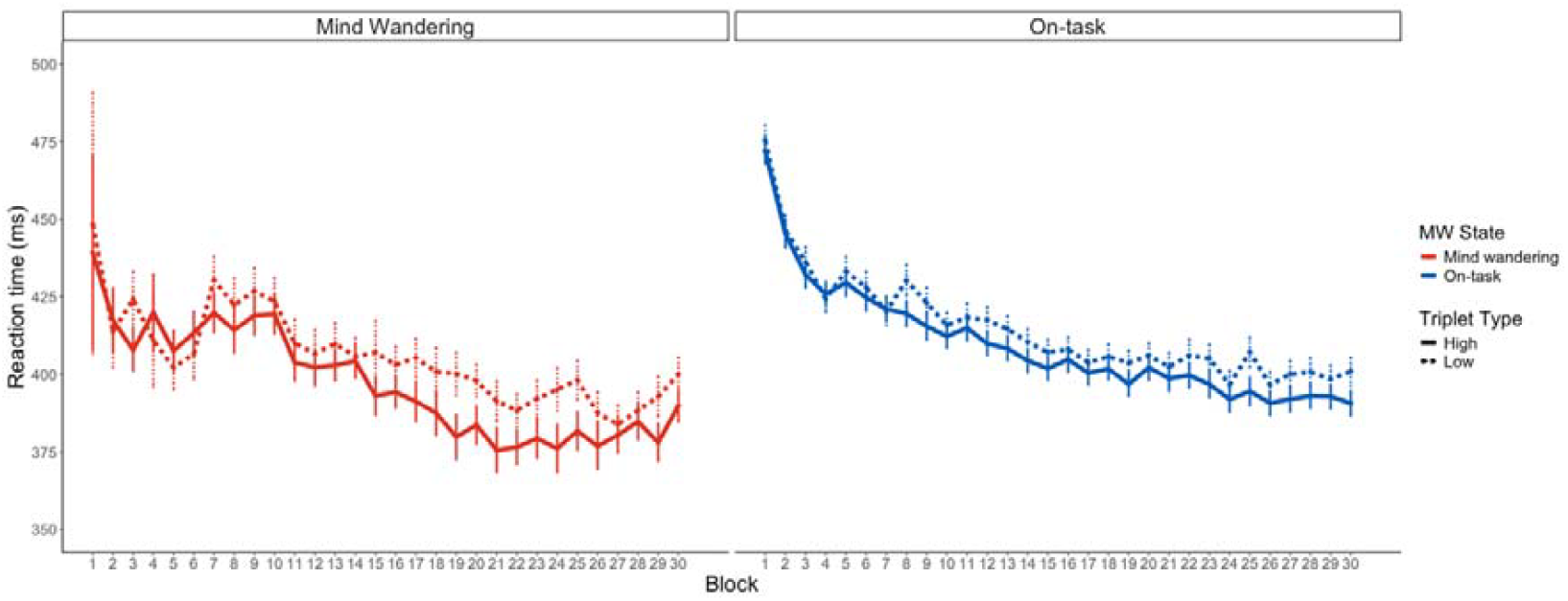
Statistical learning in RT and MW. The x-axis displays the blocks of the ASRT task, with the y-axis showing raw RTs in milliseconds. Whereas dashed lines indicate RTs to low-probability triplets, solid lines indicate RTs to high-probability triplets. Please note that the gap between the dashed and solid lines indicates statistical learning performance. Red indicates periods of MW, and blue signifies an on-task state. Error bars illustrate the standard error of the mean (SEM). Overall, statistical learning was enhanced (i.e., a bigger gap is observed between the dashed and solid lines) during periods of MW.

To sum up, when participants experienced MW, their visuomotor performance was associated with faster responses. Simultaneously, their RTs indicated improved statistical learning, meaning they were better at differentiating between less and more predictable elements.

### The role of response inhibition in the link between MW and statistical learning

Given our finding of improved statistical learning during periods of MW, our next objective was to investigate whether response inhibition modulates this relationship. We ran an LMM that included the No-Go Performance (percentage of correct ‘No-Go’ responses) as a continuous fixed effect. The dependent variable was RT on ‘Go’ trials.

The model achieved final convergence, incorporating random slopes for Block, MW, and their interaction across participants. The AIC comparison of the models without (*AIC* = 37206) and with the No-Go performance fixed effect (*AIC* = 36225) indicated better model fit for the more complex model (χ2_(8)_ = 997.104, *p* < .001). Main effects for Triplet Type and Block, as well as the interaction between MW and Block remained significant (see full results in Supplementary Material S1.4). The main effect of No-Go performance was also significant (*F*_(1, 10870.49)_ = 835.33, *p* < .001), indicating faster RT in blocks of worse No-Go Performance (*b* = 1.43, *SE* = .050). The interaction between No-Go Performance and Block was also significant (*F*_(1, 9651.35)_ = 4.05, *p* = .044). To decompose the interaction between No-Go Performance and task progression, we performed a simple slopes analysis estimating the linear trend of the Block factor at three levels of No-Go accuracy (-1 SD, mean, and +1 SD). Although RTs consistently decreased across blocks representing general visuomotor improvement (weak No-Go Performance: *b* = -0.262, *SE* = 0.026; mean No-Go Performance: *b* = -0.279, *SE* = 0.026; good No-Go Performance: *b* = -0.297, *SE* = 0.027), pairwise comparisons of these slopes revealed no statistically significant differences (all *p* > .109). A MW by No-Go Performance interaction appeared (*F*_(1, 8957.74)_ = 14.01, *p* < .001). To further elucidate the interaction, we compared the linear trends of No-Go accuracy between MW and on-task periods. The analysis revealed a significant positive relationship between No-Go performance and RT in both states; however, this slope was steeper during MW (*b* = 1.62, *SE* = 0.083) compared to on-task periods (*b* = 1.25, *SE* = 0.053). The contrast between these estimated marginal trends was statistically significant (*b* = 0.367, *SE* = 0.098, *p* < .001), indicating that the tendency for weaker inhibitory control to be associated with faster responses is significantly exacerbated when participants are in a MW state. The three-way interaction between Block, No-Go accuracy, and MW was also significant (*F*_(1, 9009.04)_ = 12.17, *p* < .001). To unpack the significant three-way interaction, we examined the linear trends of the Block factor (representing the rate of general visuomotor improvement) across MW and on-task periods at three standardized levels of No-Go Performance (-1 SD, mean, and +1 SD). When response inhibition was weak (-1 SD), RT decreased at a similar rate during both MW (*b* = -0.242, *SE* = 0.035) and on-task periods (b = -0.281, *SE* = 0.026), with no significant difference between the slopes (*b* = 0.038, *SE* = 0.032, *p* = .225). In contrast, at average and high levels of response inhibition, the general RT reduction was significantly blunted during MW compared to on-task periods. Specifically, for average No-Go performance, the slope was significantly shallower during periods of MW (*b* = -0.230, *SE* = 0.034) than during on-task periods (*b* = -0.329, *SE* = 0.023; contrast = 0.099, *SE* = 0.028, *p* < .001). This divergence was even more pronounced when response inhibition was strong (+1 SD), with the learning rate during periods of MW remaining relatively shallow (*b* = -0.217, *SE* = 0.039) compared to the steep improvement seen during on-task focus (*b* = -0.377, *SE* = 0.024; contrast = 0.160, *SE* = 0.035, *p* < .001).

Concerning statistical learning performance, a significant interaction between Triplet Type and No-Go Performance emerged (*F*_(1, 13233.76)_ = 31.52, *p* < .001). To investigate this interaction, we compared the linear trends of No-Go accuracy on RT for high-and low-probability trials. The analysis revealed a significant positive relationship between No-Go performance and RT for both trial types, but this slope was significantly steeper for high-probability trials (*b* = 1.68, *SE* = 0.066) compared to low-probability trials (*b* = 1.18, *SE* = 0.067). The contrast between these linear trends was highly significant (*b* = 0.499, *SE* = 0.089, *p* < .001), confirming that as response inhibition weakens, the divergence in RT between high-and low-probability trials becomes progressively larger. The three-way interaction between Triplet Type, No-Go Performance, and Block was also significant (*F*_(1, 13253.20)_ = 7.03, *p* = .008). To deconstruct this interaction, we compared the linear trends of RT across blocks for high-and low-probability trials at three standardized levels of No-Go Performance (-1 SD, mean, and +1 SD). The analysis revealed that the difference between high-and low-probability trials significantly increased as the task progressed only when response inhibition was weak. Specifically, for participants with weak No-Go performance (-1 SD), RT decreased at a steeper rate for high-probability trials (*b* = -0.291, *SE* = 0.029) compared to low-probability trials (*b* = -0.232, *SE* = 0.029; contrast = -0.060, *SE* = 0.021, *p* = .005). In contrast, at mean and strong (+1 SD) levels of No-Go Performance, the rate of RT improvement did not significantly differ between trial types (mean No-Go Performance contrast: *p* = .257; strong No-Go Performance contrast: *p* = .362). Importantly, a three-way interaction between Triplet Type, MW, and No-Go Performance emerged (*F*_(1, 13244.54)_ = 7.88, *p* = .005). We evaluated the estimated marginal trends to determine how the magnitude of statistical learning (the RT difference between high-and low-probability trials) changed continuously as a function of response inhibition. This trend analysis revealed a significant interaction contrast, confirming that the inverse relationship between No-Go accuracy and statistical learning was significantly modulated by the participants’ attentional state. Specifically, while poorer response inhibition was associated with a bigger difference between high-and low-probability trials during both MW (*b* = 0.749, *SE* = 0.149, *p* < .001) and on-task periods (*b* = 0.250, SE = 0.097, *p* = .010), the slope was significantly steeper during periods of MW (*b* = 0.499, *SE* = 0.178, *p* = 0.005) (Figure 5). The four-way interaction between Triplet Type, No-Go Performance, MW and Block was also significant (*F*_(1, 13259.74)_ = 5.65, *p* = .017). The interaction contrasts revealed that the difference in statistical learning trajectories between MW and on-task periods did not reach statistical significance at any level of inhibitory control. Specifically, the divergence in learning rates remained non-significant at low (*b* = -0.057, *SE* = 0.043, *p* = .181), average (*b* = 0.016, *SE* = 0.033, *p* = .636), and high (*b* = 0.089, *SE* = 0.048, *p* = .063) levels of No-Go Performance. Thus, while the overall magnitude of statistical learning is modulated by the interplay of MW and executive control, the continuous linear rate of this learning across the task does not significantly diverge between attentional states at any specific level of response inhibition. No other interactions were significant (see full results in Supplementary Material S1.4).

**Figure 5.**
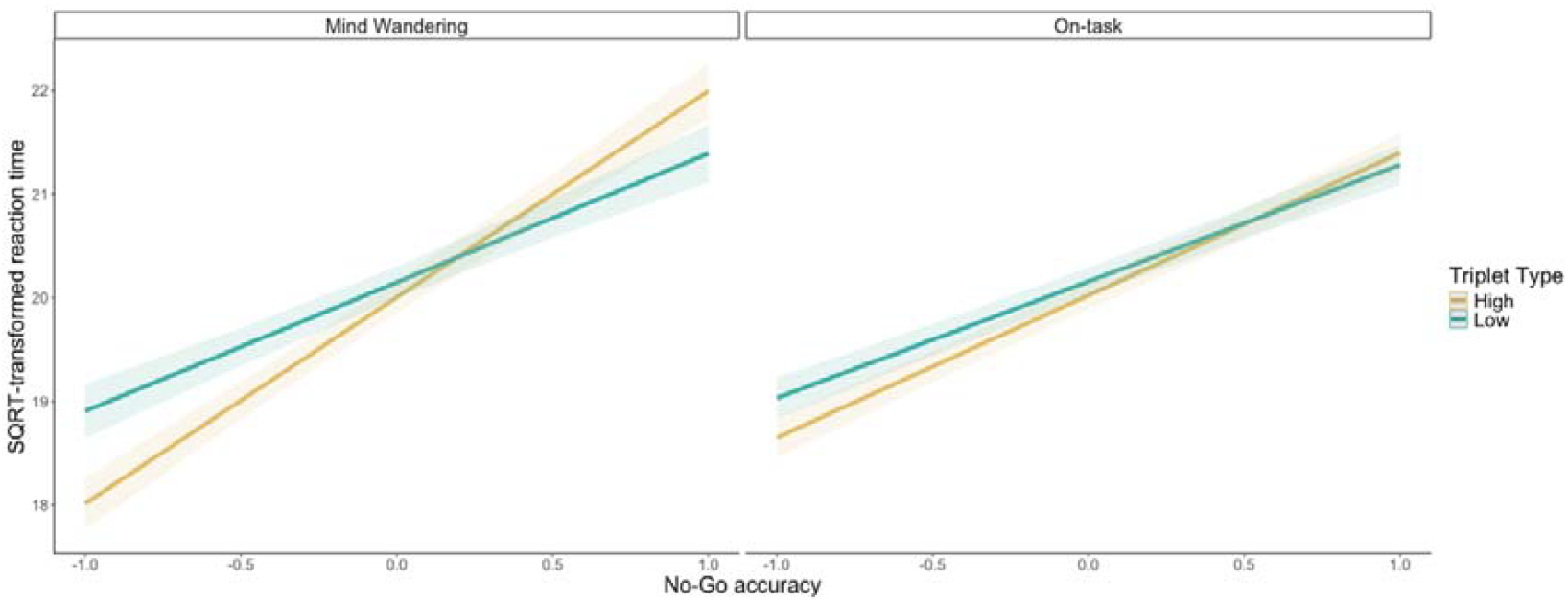
The effect of ‘No-Go’ accuracy on statistical learning in RT. The y-axis shows SQRT-transformed RT, while the x-axis represents centered ‘No-Go’ accuracy, with higher values representing better response inhibition. The green and yellow lines indicate the estimated marginal means for low-and high-probability trials, respectively. Separate subplots differentiate this relationship during MW (left) and on-task periods (right). Error bars indicate SEM. Critically, during MW, statistical learning (the difference between high-and low-probability trials) is more pronounced with weaker ‘No-Go’ accuracy.

These results demonstrate that the effect of MW on statistical learning is dependent on response inhibition. This dynamic indicates that the temporary suppression of executive control during MW promotes efficient behavioral performance and possibly the continuous refinement of statistical regularities. To sum up, MW systematically modulates the relationship between response inhibition, visuomotor performance, and the output of statistical learning. Specifically, when response inhibition is poor, MW is linked to quicker visuomotor responses and a more pronounced, automatic manifestation of learned statistical patterns.

## Discussion

This study investigated the relationship between MW, response inhibition, and statistical learning within a single experimental design. Our findings largely replicate previous research regarding the distinct effects of MW on executive control and statistical learning, while providing new insights into how these cognitive processes interact. Consistent with existing literature (Kam & Handy, 2014; Maillet et al., 2020), we observed that MW was associated with impaired response inhibition, as evidenced by decreased accuracy on ‘No-Go’ trials during MW periods. This result suggests that when attention drifts away from the task, individuals are less effective at suppressing prepotent responses. Simultaneously, previous studies (Simor et al., 2025; Vékony et al., 2025) have already shown that periods of MW resulted in impaired visuomotor performance and enhanced statistical learning. Crucially, however, our study revealed a significant modulation of the relationship between MW and statistical learning through response inhibition. Given the continuous and dynamic nature of the probabilistic task employed in our study, statistical learning here encompasses both the ongoing extraction of environmental regularities and their concurrent behavioral expression. Specifically, the findings demonstrate that when top-down control decreases, the implicit system is released from cognitive competition, leading to an optimal neurocognitive state that facilitates this ongoing, dynamic learning process.

We found that weaker response inhibition was associated with greater statistical learning during periods of MW. This suggests a potential trade-off: when executive resources are temporarily depleted due to MW, the brain may become more attuned to implicit statistical regularities (Simor et al., 2025; Vékony et al., 2025), leading to enhanced learning in terms of speed of processing. On the other hand, robust response inhibition performance was associated with diminished statistical learning during periods of MW. This aligns with the competition framework, which posits an antagonistic relationship where statistical learning thrives when cognitive control is less engaged – but now we demonstrate that this relationship is modulated by the focus of attention (MW vs. on-task). Previous research has suggested a negative association between statistical learning performance and control functions mediated by the prefrontal cortex. For instance, studies have shown negative correlations between statistical learning and frontoparietal network activity (Tóth et al., 2017), and inhibitory non-invasive brain stimulation targeting the dorsolateral prefrontal cortex has been shown to improve predictive processing as measured by statistical learning (Ambrus et al., 2020; Szücs-Bencze et al., 2025). Our direct measurements corroborate and extend these prior indications, providing empirical evidence for this inverse relationship. The observed increase in statistical learning during MW in our study is particularly salient given that MW is inherently linked to impaired task-associated cognitive control, consistent with the executive failure view (McVay & Kane, 2010). This view posits that MW episodes arise from an inability to maintain current goals, leading to reduced sustained task focus and increased susceptibility to task-unrelated interference.

Episodes of MW have also been coupled with increased activity in the frontoparietal network, providing a neural basis for how the brain’s executive system may be transiently down-regulated to facilitate the extraction of statistical regularities from the environment (Christoff et al., 2016; Smallwood et al., 2012b). Our direct behavioral measures of executive functions during MW similarly revealed a decrease in executive performance, directly supporting the notion of executive failure during these periods. This direct evidence for compromised executive control during MW, alongside enhanced statistical learning, provides compelling support for a modulatory role of executive functions in statistical learning. Our direct measurement approach provides a robust foundation for such investigations into the nuanced interplay between these fundamental cognitive processes.

Beyond the association between MW and executive control, our results also revealed significant temporal changes as the task progressed. These temporal dynamics can be understood through the lens of the vigilance decrement literature, particularly the Resource-Control Account of sustained attention (Thomson et al., 2015). According to this framework, maintaining on-task attention requires the continuous exertion of executive control that inevitably depletes over time. Consistent with recent empirical findings that demonstrate a simultaneous decline in cognitive control and sustained attention across time on task (Luna et al., 2022), our results show a temporal reduction in response inhibition accompanied by an increase in MW. Expanding upon the Resource-Control Account, our findings suggest that this time-dependent decrease in executive control and the consequent rise in MW do not merely represent a cognitive failure. Rather, this progressive executive depletion mediates a shift in the neurocognitive state that facilitates statistical learning.

The enhanced difference between high-and low-probability trials during MW offers two complementary theoretical interpretations: a learning-based account, in which MW actively modulates the ongoing acquisition of statistical regularities, and a performance-based account, in which MW primarily enhances the behavioral expression of previously acquired statistical knowledge. The acquisition and expression are dynamically intertwined in continuous probabilistic tasks; MW might primarily drive the learning process itself. By withdrawing effortful, rule-based attention from the environment, MW creates a neurocognitive state that shields the implicit system from interference, thereby facilitating not just the execution but also the continuous extraction and updating of complex statistical regularities (Decker et al., 2022; Tan et al., 2015). According to the alternative, performance-based interpretation, aligning with the distinction between learning and performance (Soderstrom & Bjork, 2015; Vékony et al., 2020, 2022), the attenuation of top-down executive control during MW may temporarily release the implicit system from cognitive competition, allowing participants to exhibit more automatic, learned behavior (Jiménez et al., 2009; Prutean et al., 2022; Vaquero et al., 2019). This framework accounts for the observed interactions with response inhibition: when response inhibition is low, automated routines take over. However, the interaction with attentional state suggests that failures in response inhibition during MW reflect a systemic shift that optimizes the implicit system for learning. During MW, a No-Go error reflects a systemic withdrawal of top-down constraints, leaving automated routines to run completely unchecked. In contrast, during on-task periods, No-Go errors likely reflect transient resource depletion or impulsivity while the controlled system is still actively, albeit unsuccessfully, attempting to supervise behavior. However, temporal dynamics reported by Vékony et al. (2025) without a demanding response inhibition component demonstrated that the MW-related enhancement of statistical learning progressively diminished as the task advanced, suggesting that the learning component plays a more prominent role in driving the observed effects. We acknowledge that both mechanisms could contribute to the adaptive advantage of transient executive lapses for implicit cognitive processing. Future research is warranted to clarify the specific role these two mechanisms play in the observed effects.

We speculate that during offline moments such as MW, the brain replays previously encountered sequences and engages in internal simulations of potential sequence variations and transitions. This generative process may allow the brain to explore alternative combinations of events beyond those explicitly experienced, effectively filling in gaps and refining its internal model of the underlying structure. Such simulations likely support the abstraction of statistical regularities by amplifying co-occurrence patterns and reinforcing predictive associations, even in the absence of direct sensory input. In this way, transient offline states such as MW may serve as a window for unsupervised internal modeling, contributing to more efficient statistical learning. Our findings, together with previous research (Simor et al., 2025; Vékony et al., 2025), raise the possibility that this process may be driven by memory consolidation associated with local sleep occurring in the waking brain (Wamsley, 2022; Wamsley & Summer, 2020). Statistical learning was most pronounced when inhibitory control was at its lowest, aligning with theories suggesting that local-sleep-like neural states may enhance model-free learning by attenuating frontoparietal executive networks. This interpretation is consistent with the idea that local-sleep-dependent consolidation stabilizes recent memories and supports the formation of predictive models of environmental regularities (Vékony et al., 2025). However, more studies using methods like magnetoencephalography or intracranial electroencephalography are needed to explore the link between MW, local sleep activity, and how we develop predictive models.

Human cognition must balance the demands of goal-directed focus with the need to learn from the environment. Here, we dissect this balance and reveal a key mechanism governing the interplay between internal thoughts and statistical learning. We demonstrate that the enhancement of statistical learning during MW is directly modulated by the efficacy of response inhibition. While we confirmed previous findings that MW diminishes inhibitory control and accelerates responses to predictable patterns, our significant advancement is demonstrating that these effects are not parallel. The magnitude of the learning benefit during MW relies on the inadequacy of inhibitory control, providing evidence for this functional trade-off. Whether this executive dysfunction primarily facilitates the ongoing acquisition or the concurrent behavioral expression of implicit learning remains a question for future research. This inverse relationship offers support for neurocompetition models that posit an antagonistic relationship between executive and implicit statistical learning systems. These results reframe the adaptive function of MW, suggesting that it may represent a cognitive state optimized not for immediate goal pursuit but for the continuous, unsupervised acquisition of a predictive model. This dynamic interplay between internally-and externally-directed cognitive modes appears to be a fundamental mechanism that allows for flexible adaptation in a complex and ever-changing environment.

## Funding

This work was supported by the the French National Grant Agency (ANR-24-CE37-5807); the National Brain Research Program project NAP2022-I-2/2022 of the Hungarian Academy of Sciences (D.N.); the Hungarian National Research, Development and Innovation Office grant NKFI FK 142945 (P.S.) and 128016 (D.N.); the Spanish Ministry of Science, Innovation and Universities (MICIU), the State Research Agency (AEI), and the European Regional Development Fund (FEDER, UE) through the grant PID2024-160183NA-I00 (MICIU/AEI/10.13039/501100011033/FEDER, UE) (T.V.); and the Fonds de la Recherche Scientifique – FNRS (F.R.S.-FNRS) through a 2025 Mandat d’Impulsion Scientifique (MIS 40036982) (P.S.). PS is a research associate at the F.R.S.-FNRS.

## Author contributions

Conceptualization: T.V., G.C., P.S., and D.N.; methodology: T.V., B.B., G.C., P.S., and D.N.; software: T.V.; validation: T.V., G.C., P.S., and D.N.; formal analysis: T.V. and B.B.; investigation: T.V. and D.N.; resources: T.V., B.B., G.C., P.S., and D.N.; data curation: T.V.; writing—original draft: T.V., G.C., P.S., and D.N.; writing—review and editing: T.V., B.B., G.C., P.S., and D.N.; visualization: T.V.; supervision: G.C., P.S., and D.N.; project administration: T.V., G.C., P.S., and D.N.; funding acquisition: P.S., D.N and T.V.

## Declaration of interests

The authors declare no competing interests.

## Data availability statement

The raw data and all codes required to duplicate the results, figures, and tables are accessible at the following OSF repository: https://osf.io/53jqs/?view_only=37ef7f12c7c3416fbf9992b6c4109624

## Notes

### Competing Interest Statement

The authors have declared no competing interest.

### Summary of Updates

Updated version after revision (additions to the introduction and discussion sections, a change in the post-hoc strategy in the results section, and methodological clarifications, updated affiliations and contact information)

## References

Ambrus, G. G., Vékony, T., Janacsek, K., Trimborn, A. B. C., Kovács, G., & Nemeth, D. (2020). When less is more: Enhanced statistical learning of non-adjacent dependencies after disruption of bilateral DLPFC. Journal of Memory and Language, 114, 104144. 10.1016/j.jml.2020.104144

Amer, T., Campbell, K. L., & Hasher, L. (2016). Cognitive Control As a Double-Edged Sword. Trends in Cognitive Sciences, 20(12), 905–915. 10.1016/J.TICS.2016.10.002

Andrillon, T., Burns, A., Mackay, T., Windt, J., & Tsuchiya, N. (2021). Predicting lapses of attention with sleep-like slow waves. Nature Communications, 12(1), 1–12. 10.1038/s41467-021-23890-7

Arnau, S., Löffler, C., Rummel, J., Hagemann, D., Wascher, E., & Schubert, A. L. (2020). Inter-trial alpha power indicates mind wandering. Psychophysiology, 57(6), e13581. 10.1111/PSYP.13581

Bastian, M., & Sackur, J. (2013). Mind wandering at the fingertips: Automatic parsing of subjective states based on response time variability. Frontiers in Psychology, 4(SEP), 573. 10.3389/fpsyg.2013.00573

Boayue, N. M., Csifcsák, G., Kreis, I. V., Schmidt, C., Finn, I., Hovde Vollsund, A., & Mittner, M. E. (2021). The interplay between executive control, behavioural variability and mind wandering: Insights from a high-definition transcranial direct-current stimulation study. European Journal of Neuroscience, 53(5), 1498–1516. 10.1111/EJN.15049

Cheyne, J. A., Solman, G. J. F., Carriere, J. S. A., & Smilek, D. (2009). Anatomy of an error: A bidirectional state model of task engagement/disengagement and attention-related errors. Cognition, 111(1), 98–113. 10.1016/j.cognition.2008.12.009

Christoff, K., Irving, Z. C., Fox, K. C. R., Spreng, R. N., & Andrews-Hanna, J. R. (2016). Mind-wandering as spontaneous thought: A dynamic framework. Nature Reviews Neuroscience, 17(11), 718–731. 10.1038/nrn.2016.113

de Leeuw, J. R. (2015). jsPsych: A JavaScript library for creating behavioral experiments in a Web browser. Behavior Research Methods, 47(1), 1–12. 10.3758/s13428-014-0458-y

Decker, A., Dubois, M., Duncan, K., & Finn, A. S. (2022). Pay attention and you might miss it: Greater learning during attentional lapses. Psychonomic Bulletin & Review 2022 30:3, 30(3), 1041–1052. 10.3758/S13423-022-02226-6

Dong, H. W., Mills, C., Knight, R. T., & Kam, J. W. Y. (2021). Detection of mind wandering using EEG: Within and across individuals. PLOS ONE, 16(5), e0251490. 10.1371/JOURNAL.PONE.0251490

Howard, J. H., & Howard, D. V. (1997). Age differences in implicit learning of higher order dependencies in serial patterns. Psychology and Aging, 12(4), 634–656. 10.1037/0882-7974.12.4.634

Jiménez, L., Lupiáñez, J., & Vaquero, J. M. M. (2009). Sequential congruency effects in implicit sequence learning. Consciousness and Cognition, 18(3), 690–700. 10.1016/J.CONCOG.2009.04.006

Kam, J. W. Y., Dao, E., Farley, J., Fitzpatrick, K., Smallwood, J., Schooler, J. W., & Handy, T. C. (2011). Slow Fluctuations in Attentional Control of Sensory Cortex. Journal of Cognitive Neuroscience, 23(2), 460–470. 10.1162/JOCN.2010.21443

Kam, J. W. Y., & Handy, T. C. (2014). Differential recruitment of executive resources during mind wandering. Consciousness and Cognition, 26(1), 51–63. 10.1016/J.CONCOG.2014.03.002

Kam, J. W. Y., Handy, T. C., Smallwood, J., & Allen, M. (2013). The neurocognitive consequences of the wandering mind: a mechanistic account of sensory-motor decoupling. Frontiers in Psychology, 4, 55071. 10.3389/FPSYG.2013.00725

Liu, S., Rabovsky, M., & Schad, D. J. (2023). Spontaneous mind wandering impairs model-based decision making. PLoS ONE, 18(1 January), e0279532. 10.1371/journal.pone.0279532

Luna, F. G., Tortajada, M., Martín-Arévalo, E., Botta, F., & Lupiáñez, J. (2022). A vigilance decrement comes along with an executive control decrement: Testing the resource-control theory. Psychonomic Bulletin & Review 2022 29:5, 29(5), 1831–1843. 10.3758/S13423-022-02089-X

Maillet, D., Yu, L., Hasher, L., & Grady, C. L. (2020). Age-related differences in the impact of mind-wandering and visual distraction on performance in a go/no-go task. Psychology and Aging, 35(5), 627–638. 10.1037/PAG0000409

McVay, J. C., & Kane, M. J. (2010). Does mind wandering reflect executive function or executive failure? Comment on Smallwood and Schooler (2006) and Watkins (2008). Psychological Bulletin, 136(2), 188–197. 10.1037/a0018298

McVay, J. C., & Kane, M. J. (2012). Why does working memory capacity predict variation in reading comprehension? On the influence of mind wandering and executive attention. Journal of Experimental Psychology: General, 141(2), 302–320. 10.1037/a0025250

Meteyard, L., & Davies, R. A. I. (2020). Best practice guidance for linear mixed-effects models in psychological science. Journal of Memory and Language, 112, 104092. 10.1016/J.JML.2020.104092

Mooneyham, B. W., & Schooler, J. W. (2013). The costs and benefits of mind-wandering: A review. Canadian Journal of Experimental Psychology, 67(1), 11–18. 10.1037/a0031569

Mrazek, M. D., Smallwood, J., Franklin, M. S., Chin, J. M., Baird, B., & Schooler, J. W. (2012). The role of mind-wandering in measurements of general aptitude. Journal of Experimental Psychology: General, 141(4), 788–798. 10.1037/a0027968

Mrazek, M. D., Smallwood, J., & Schooler, J. W. (2012). Mindfulness and mind-wandering: Finding convergence through opposing constructs. Emotion, 12(3), 442–448. 10.1037/a0026678

Nemeth, D., Janacsek, K., Polner, B., & Kovacs, Z. A. (2013). Boosting human learning by hypnosis. Cerebral Cortex, 23(4), 801–805. 10.1093/cercor/bhs068

Prutean, N., Wenk, T., Leiva, A., Vaquero, J. M. M., Lupiáñez, J., & Jiménez, L. (2022). Cognitive Control Modulates the Expression of Implicit Sequence Learning: Congruency Sequence and Oddball-Dependent Sequence Effects. Journal of Experimental Psychology: Human Perception and Performance, 48(8), 842–855. 10.1037/XHP0001025

Randall, J. G., Oswald, F. L., & Beier, M. E. (2014). Mind-wandering, cognition, and performance: a theory-driven meta-analysis of attention regulation. Psychological Bulletin, 140(6), 1411–1431. 10.1037/A0037428.

Rodd, J. M. (2024). Moving experimental psychology online: How to obtain high quality data when we can’t see our participants. Journal of Memory and Language, 134, 104472. 10.1016/J.JML.2023.104472

Schooler, J. W., Smallwood, J., Christoff, K., Handy, T. C., Reichle, E. D., & Sayette, M. A. (2011). Meta-awareness, perceptual decoupling and the wandering mind. Trends in Cognitive Sciences, 15(7), 319–326. 10.1016/j.tics.2011.05.006

Seli, P., Smallwood, J., Cheyne, J. A., & Smilek, D. (2015). On the relation of mind wandering and ADHD symptomatology. Psychonomic Bulletin and Review, 22(3), 629–636. 10.3758/s13423-014-0793-0

Simor, P., Vékony, T., Farkas, B. C., Szalárdy, O., Bogdány, T., Brezóczki, B., Csifcsák, G., & Németh, D. (2025). Mind Wandering during Implicit Learning Is Associated with Increased Periodic EEG Activity and Improved Extraction of Hidden Probabilistic Patterns. Journal of Neuroscience, 45(19). 10.1523/JNEUROSCI.1421-24.2025

Smalle, E. H. M., Panouilleres, M., Szmalec, A., & Möttönen, R. (2017). Language learning in the adult brain: Disrupting the dorsolateral prefrontal cortex facilitates word-form learning. Scientific Reports, 7(1), 13966. 10.1038/s41598-017-14547-x

Smallwood, J., Brown, K., Baird, B., & Schooler, J. W. (2012a). Cooperation between the default mode network and the frontal–parietal network in the production of an internal train of thought. Brain Research, 1428, 60–70. 10.1016/J.BRAINRES.2011.03.072

Smallwood, J., Brown, K., Baird, B., & Schooler, J. W. (2012b). Cooperation between the default mode network and the frontal–parietal network in the production of an internal train of thought. Brain Research, 1428, 60–70. 10.1016/J.BRAINRES.2011.03.072

Smallwood, J., & Schooler, J. W. (2006). The restless mind. Psychological Bulletin, 132(6), 946–958. 10.1037/0033-2909.132.6.946

Soderstrom, N. C., & Bjork, R. A. (2015). Learning versus performance: An integrative review. Perspectives on Psychological Science, 10(2), 176–199. 10.1177/1745691615569000

Song, S., Howard, J. H., & Howard, D. V. (2007). Implicit probabilistic sequence learning is independent of explicit awareness. Learning and Memory, 14(3), 167–176. 10.1101/lm.437407

Szücs-Bencze, L., Vékony, T., Pesthy, O., Kocsis, K., Kincses, Z. T., Szabó, N., & Nemeth, D. (2025). Enhancing retrieval capacity of the predictive brain through dorsolateral prefrontal cortex intervention. Cerebral Cortex, 35(2). 10.1093/CERCOR/BHAF005

Tan, T., Zou, H., Chen, C., & Luo, J. (2015). Mind Wandering and the Incubation Effect in Insight Problem Solving. Creativity Research Journal, 27(4), 375–382. 10.1080/10400419.2015.1088290

Thompson-Schill, S. L., Ramscar, M., & Chrysikou, E. G. (2009). Cognition without control: When a little frontal lobe goes a long way. Current Directions in Psychological Science, 18(5), 259–263. 10.1111/j.1467-8721.2009.01648.x

Thomson, D. R., Besner, D., & Smilek, D. (2015). A Resource-Control Account of Sustained Attention: Evidence From Mind-Wandering and Vigilance Paradigms. Perspectives on Psychological Science, 10(1), 82–96. 10.1177/1745691614556681

Tóth, B., Janacsek, K., Takács, Á., Kóbor, A., Zavecz, Z., & Nemeth, D. (2017). Dynamics of EEG functional connectivity during statistical learning. Neurobiology of Learning and Memory, 144, 216–229. 10.1016/j.nlm.2017.07.015

Vaquero, J. M. M., Jiménez, L., & Lupiáñez, J. (2006). The problem of reversals in assessing implicit sequence learning with serial reaction time tasks. Experimental Brain Research 2006 175:1, 175(1), 97–109. 10.1007/S00221-006-0523-6

Vaquero, J. M. M., Lupiáñez, J., & Jiménez, L. (2019). Asymmetrical effects of control on the expression of implicit sequence learning. Psychological Research 2019 84:8, 84(8), 2157–2171. 10.1007/S00426-019-01222-1

Vékony, T., Farkas, B. C., Brezóczki, B., Mittner, M., Csifcsák, G., Simor, P., & Németh, D. (2025). Mind wandering enhances statistical learning. IScience, 111703. 10.1016/J.ISCI.2024.111703

Vékony, T., Marossy, H., Must, A., Vécsei, L., Janacsek, K., & Nemeth, D. (2020). Speed or accuracy instructions during skill learning do not affect the acquired knowledge. Cerebral Cortex Communications, 1(1). 10.1093/texcom/tgaa041

Vékony, T., Pleche, C., Pesthy, O., Janacsek, K., & Nemeth, D. (2022). Speed and accuracy instructions affect two aspects of skill learning differently. Npj Science of Learning, 7(1), 1–11. 10.1038/s41539-022-00144-9

Verschooren, S., & Egner, T. (2023). When the mind’s eye prevails: The Internal Dominance over External Attention (IDEA) hypothesis. Psychonomic Bulletin & Review 2023 30:5, 30(5), 1668– 1688. 10.3758/S13423-023-02272-8

Virag, M., Janacsek, K., Horvath, A., Bujdoso, Z., Fabo, D., & Nemeth, D. (2015). Competition between frontal lobe functions and implicit sequence learning: evidence from the long-term effects of alcohol. Experimental Brain Research, 233(7), 2081–2089. 10.1007/s00221-015-4279-8

Wamsley, E. J. (2022). Offline memory consolidation during waking rest. Nature Reviews Psychology, 1(8), 441–453. 10.1038/s44159-022-00072-w

Wamsley, E. J., & Summer, T. (2020). Spontaneous entry into an “offline” state during wakefulness: A mechanism of memory consolidation? Journal of Cognitive Neuroscience, 32(9), 1714–1734. 10.1162/jocn_a_01587

