## Supplementary Material for "A functional trade-off between executive control and implicit statistical learning is dynamically gated by mind wandering"

**Supplementary Materials**

**S1. Details of the models presented in the main text**

**S1.1 Response inhibition and MW**

**Table S1.**

| *Terms* | *b* | *SE b* | *95% CI* | *t* | *df* | *p* |
| --- | --- | --- | --- | --- | --- | --- |
| MW [On-task] | -0.0461 | 0.0038 | -0.0537 – -0.0385 | -11.9836 | 183.0220 | **<0.001** |
| Block | -0.0526 | 0.0043 | -0.0610 – -0.0442 | -12.3364 | 212.9757 | **<0.001** |
| MW [On-task] × Block | -0.0011 | 0.0028 | -0.0067 – 0.0045 | -0.3743 | 155.6834 | 0.709 |
| **Random Effects** |  | | | | | |
| σ^2^ | 0.02 | | |  | | |
| τ_00_ _ID_ | 0.02 | | |  | | |
| τ_11_ _ID.Block_ | 0.00 | | |  | | |
| τ_11_ _ID.MW [On-task]_ | 0.00 | | |  | | |
| τ_11_ _ID.Block:MW [On-task]_ | 0.00 | | |  | | |
| ρ_01_ | 0.08 | | |  | | |
|  | 0.42 | | |  | | |
|  | -0.02 | | |  | | |
| ICC | 0.54 | | |  | | |
| N _ID_ | 240 | | |  | | |
| Observations | 14340 | | |  | | |
| Marginal R^2^ / Conditional R^2^ | 0.098 / 0.582 | | |  | | |

*Note.* Response inhibition became weaker as the task progressed (main effect of Block), and it was generally weaker during periods of MW (main effect of MW).

**S1.2 Statistical learning and visuomotor performance**

**Table S2.**

| *Terms* | *b* | *SE b* | *95% CI* | *t* | *df* | *p* |
| --- | --- | --- | --- | --- | --- | --- |
| (Intercept) | 20.0963 | 0.0689 | 19.9606 – 20.2320 | 291.7479 | 238.9964 | **<0.001** |
| Triplet Type [Low] | -0.0786 | 0.0075 | -0.0933 – -0.0638 | -10.5088 | 227.8849 | **<0.001** |
| Block | -0.4020 | 0.0279 | -0.4569 – -0.3471 | -14.4313 | 236.6228 | **<0.001** |
| Triplet Type [Low] × Block | -0.0294 | 0.0071 | -0.0432 – -0.0156 | -4.1663 | 13626.0434 | **<0.001** |
| **Random Effects** | | | | | | |
| σ^2^ | 0.71 | | | | | |
| τ_00_ _ID_ | 1.13 | | | | | |
| τ_11_ _ID.Block_ | 0.17 | | | | | |
| τ_11_ _ID.Triplet Type [Low]_ | 0.00 | | | | | |
| ρ_01_ | -0.13 | | | | | |
|  | 0.28 | | | | | |
| N _ID_ | 0.65 | | | | | |
| Observations | 240 | | | | | |
| Marginal R^2^ / Conditional R^2^ | 14340 | | | | | |

*Note.* Statistical learning was demonstrated as RTs for high-probability triplets were generally faster than those for low-probability triplets (main effect of Triplet Type). Reaction times improved as the task advanced (main effect of Block). The interplay between Triplet Type and Block demonstrated more significant statistical learning (the difference between high- and low-probability triplets) as the task progressed. **S1.3 Statistical learning, visuomotor performance, and MW**

**Table S3.**

| *Terms* | *b* | *SE b* | *95% CI* | *t* | *df* | *p* |
| --- | --- | --- | --- | --- | --- | --- |
| (Intercept) | 20.0530 | 0.0700 | 19.9150 – 20.1909 | 286.5098 | 228.7006 | **<0.001** |
| Triplet Type [Low] | -0.0841 | 0.0079 | -0.0995 – -0.0686 | -10.6669 | 13246.6069 | **<0.001** |
| MW [on-task] | -0.0841 | 0.0239 | -0.1315 – -0.0367 | -3.5166 | 113.2607 | **0.001** |
| Block | -0.3617 | 0.0281 | -0.4170 – -0.3064 | -12.8923 | 215.0626 | **<0.001** |
| Triplet Type [Low] × MW [on-task] | -0.0189 | 0.0079 | -0.0344 – -0.0035 | -2.3983 | 13247.7292 | **0.016** |
| Triplet Type [Low] × Block | -0.0311 | 0.0081 | -0.0469 – -0.0153 | -3.8520 | 13249.2109 | **<0.001** |
| MW [on-task] × Block | 0.0438 | 0.0150 | 0.0141 – 0.0735 | 2.9161 | 139.7544 | **0.004** |
| (Triplet Type [Low] × MW [on-task]) × Block | -0.0115 | 0.0081 | -0.0273 – 0.0044 | -1.4201 | 13250.7431 | 0.156 |
| **Random Effects** | | | | | | |
| σ^2^ | 0.67 | | | | | |
| τ_00_ _ID_ | 1.14 | | | | | |
| τ_11_ _ID.Block_ | 0.16 | | | | | |
| τ_11_ _ID.MW [on-task]_ | 0.10 | | | | | |
| τ_11_ _ID.Block:MW [on-task]_ | 0.02 | | | | | |
| ρ_01_ | -0.15 | | | | | |
|  | 0.23 | | | | | |
|  | 0.16 | | | | | |
| ICC | 0.67 | | | | | |
| N _ID_ | 240 | | | | | |
| Observations | 14340 | | | | | |
| Marginal R^2^ / Conditional R^2^ | 0.079 / 0.693 | | | | | |

*Note.* RTs during periods of MW were faster compared to on-task periods (main effect of MW), which became progressively larger (interaction between MW and Block). Statistical learning (i.e., the difference between high- and low-probability trials) was overall larger during periods of MW compared to on-task periods (interaction between Triplet Type and MW).

**S1.4 The interplay of statistical learning, response inhibition, and MW**

**Table S4.**

| *Terms* | *b* | *SE b* | *95% CI* | *t* | *df* | *p* |
| --- | --- | --- | --- | --- | --- | --- |
| (Intercept) | 20.0825 | 0.0696 | 19.9454 – 20.2197 | 288.4172 | 231.8465 | **<0.001** |
| Triplet Type [Low] | -0.0693 | 0.0080 | -0.0850 – -0.0536 | -8.6674 | 13225.7116 | **<0.001** |
| No-Go | 1.4335 | 0.0496 | 1.3363 – 1.5307 | 28.9021 | 10870.4901 | **<0.001** |
| MW [on-task] | -0.0067 | 0.0207 | -0.0478 – 0.0344 | -0.3229 | 105.5973 | 0.747 |
| Block | -0.2792 | 0.0255 | -0.3294 – -0.2289 | -10.9561 | 215.6176 | **<0.001** |
| Triplet Type [Low] × No-Go | 0.2495 | 0.0444 | 0.1624 – 0.3367 | 5.6139 | 13233.7634 | **<0.001** |
| Triplet Type [Low] × MW [on-task] | -0.0025 | 0.0080 | -0.0182 – 0.0132 | -0.3097 | 13225.6529 | 0.757 |
| No-Go × MW [on-task] | 0.1837 | 0.0491 | 0.0875 – 0.2799 | 3.7430 | 8957.7425 | **<0.001** |
| Triplet Type [Low] × block | -0.0095 | 0.0084 | -0.0258 – 0.0069 | -1.1339 | 13225.6767 | 0.257 |
| No-Go × Block | -0.0981 | 0.0488 | -0.1937 – -0.0025 | -2.0117 | 9651.3506 | **0.044** |
| MW [on-task] × Block | 0.0496 | 0.0140 | 0.0219 – 0.0773 | 3.5369 | 152.4562 | **0.001** |
| (Triplet Type [Low] × No-Go) × MW [on-task] | 0.1247 | 0.0444 | 0.0376 – 0.2119 | 2.8068 | 13244.5399 | **0.005** |
| (Triplet Type [Low] × No-Go) × Block | 0.1135 | 0.0428 | 0.0296 – 0.1974 | 2.6508 | 13253.1953 | **0.008** |
| (Triplet Type [Low] × MW [on-task]) × Block | 0.0040 | 0.0084 | -0.0124 – 0.0203 | 0.4736 | 13224.9386 | 0.636 |
| (No-Go × MW [on-task] × Block | 0.1698 | 0.0487 | 0.0744 – 0.2652 | 3.4880 | 9009.0385 | **<0.001** |
| (Triplet Type [Low] × No-Go × MW [on-task] × Block | 0.1018 | 0.0428 | 0.0179 – 0.1857 | 2.3780 | 13259.7375 | **0.017** |
| **Random Effects** | | | | | | |
| σ^2^ | 0.63 | | | | | |
| τ_00_ _ID_ | 1.13 | | | | | |
| τ_11_ _ID.Block_ | 0.13 | | | | | |
| τ_11_ _ID.MW [on-task]_ | 0.07 | | | | | |
| τ_11_ _ID.Block:_ _MW [on-task]_ | 0.02 | | | | | |
| ρ_01_ | -0.21 | | | | | |
|  | 0.18 | | | | | |
|  | 0.15 | | | | | |
| ICC | 0.67 | | | | | |
| N _ID_ | 240 | | | | | |
| Observations | 14340 | | | | | |
| Marginal R^2^ / Conditional R^2^ | 0.107 / 0.708 | | | | | |

*Note.* Worse No-Go performance was associated with faster RTs (main effect of No-Go). Participants exhibited faster RTs during MW when their No-Go performance was weak, but slower RTs during MW when their No-Go performance was good (interaction between No-Go and MW). Statistical learning was better during periods of weak No-Go performance (interaction between No-Go and Triplet Type). When No-Go performance was weak, participants showed greater statistical learning during MW compared to on-task periods (interaction between No-Go, Triplet Type, and MW).

**S2. Accuracy analysis**

**S2.1 Statistical learning and visuomotor performance – Accuracy**

**Table S5.**

| *Terms* | *b* | *SE b* | *95% CI* | *t* | *df* | *p* |
| --- | --- | --- | --- | --- | --- | --- |
| (Intercept) | 0.9358 | 0.0029 | 0.9301 – 0.9416 | 321.1150 | 238.2028 | **<0.001** |
| Triplet Type [Low] | 0.0044 | 0.0009 | 0.0028 – 0.0061 | 5.1995 | 228.0869 | **<0.001** |
| Block | -0.0113 | 0.0019 | -0.0150 – -0.0077 | -6.0619 | 234.5906 | **<0.001** |
| Triplet Type [Low] × Block | 0.0001 | 0.0007 | -0.0013 – 0.0014 | 0.0724 | 13629.6022 | 0.942 |
| **Random Effects** | | | | | | |
| σ^2^ | 0.01 | | | | | |
| τ_00_ _ID_ | 0.00 | | | | | |
| τ_11_ _ID.Block_ | 0.00 | | | | | |
| τ_11_ _ID.Triplet Type [Low]_ | 0.00 | | | | | |
| ρ_01_ | 0.76 | | | | | |
|  | 0.27 | | | | | |
| ICC | 0.28 | | | | | |
| N _ID_ | 240 | | | | | |
| Observations | 14340 | | | | | |
| Marginal R^2^ / Conditional R^2^ | 0.015 / 0.295 | | | | | |

*Note.* Statistical learning was demonstrated as accuracy for high-probability triplets was generally better than those for low-probability triplets (main effect of Triplet Type). Accuracy decreased as the task advanced (main effect of Block). The interplay between Triplet Type and Block demonstrated more significant statistical learning (the difference between high- and low-probability triplets) as the task progressed.

**S2.2 Statistical learning, visuomotor performance, and MW – Accuracy**

**Table S6.**

| *Terms* | *b* | *SE b* | *95% CI* | *t* | *df* | *p* |
| --- | --- | --- | --- | --- | --- | --- |
| (Intercept) | 0.9312 | 0.0032 | 0.9248 – 0.9376 | 287.6789 | 242.2933 | **<0.001** |
| Triplet Type [Low] | 0.0044 | 0.0008 | 0.0028 – 0.0059 | 5.5577 | 13608.4702 | **<0.001** |
| MW [on-task] | -0.0127 | 0.0019 | -0.0165 – -0.0089 | -6.5657 | 180.1267 | **<0.001** |
| Block | -0.0086 | 0.0015 | -0.0115 – -0.0056 | -5.7275 | 260.4812 | **<0.001** |
| Triplet Type [Low] × MW [on-task] | -0.0009 | 0.0008 | -0.0025 – 0.0006 | -1.1811 | 13609.7025 | 0.238 |
| Triplet Type [Low] × Block | -0.0004 | 0.0008 | -0.0020 – 0.0012 | -0.4833 | 13610.2430 | 0.629 |
| MW [on-task] × Block | -0.0007 | 0.0009 | -0.0025 – 0.0012 | -0.7220 | 6813.2651 | 0.470 |
| (Triplet Type [Low] × MW [on-task]) × Block | -0.0013 | 0.0008 | -0.0029 – 0.0002 | -1.6653 | 13609.4989 | 0.096 |
| **Random Effects** | | | | | | |
| σ^2^ | 0.01 | | | | | |
| τ_00_ _ID_ | 0.00 | | | | | |
| τ_11_ _ID.Block_ | 0.00 | | | | | |
| τ_11_ _ID.MW [on-task]_ | 0.00 | | | | | |
| ρ_01_ | 0.62 | | | | | |
|  | 0.79 | | | | | |
| ICC | 0.27 | | | | | |
| N _ID_ | 240 | | | | | |
| Observations | 14340 | | | | | |
| Marginal R^2^ / Conditional R^2^ | 0.028 / 0.293 | | | | | |

*Note.* Participants were less accurate during MW compared to on-task periods (main effect of MW).

**S2.3 The interplay of statistical learning, response inhibition, and MW – Accuracy**

**Table S7.**

| *Terms* | *b* | *SE b* | *95% CI* | *t* | *df* | *p* |
| --- | --- | --- | --- | --- | --- | --- |
| (Intercept) | 0.9356 | 0.0031 | 0.9295 – 0.9416 | 304.5096 | 246.8589 | **<0.001** |
| Triplet Type [Low] | 0.0056 | 0.0010 | 0.0037 – 0.0074 | 5.8153 | 360.7345 | **<0.001** |
| No-Go | 0.1053 | 0.0048 | 0.0960 – 0.1147 | 22.0723 | 11921.5727 | **<0.001** |
| MW [on-task] | -0.0072 | 0.0017 | -0.0106 – -0.0038 | -4.1761 | 191.9529 | **<0.001** |
| Block | -0.0019 | 0.0013 | -0.0046 – 0.0007 | -1.4133 | 296.1810 | 0.159 |
| Triplet Type [Low] × No-Go | 0.0271 | 0.0045 | 0.0184 – 0.0359 | 6.0683 | 13605.0537 | **<0.001** |
| Triplet Type [Low] × MW [on-task] | 0.0008 | 0.0008 | -0.0009 – 0.0024 | 0.9159 | 4706.7899 | 0.360 |
| No-Go × MW [on-task] | 0.0301 | 0.0047 | 0.0208 – 0.0394 | 6.3548 | 13160.6206 | **<0.001** |
| Triplet Type [Low] × Block | 0.0011 | 0.0008 | -0.0005 – 0.0027 | 1.3073 | 13598.4540 | 0.191 |
| No-Go × Block | 0.0176 | 0.0047 | 0.0084 – 0.0268 | 3.7577 | 13334.2038 | **<0.001** |
| MW [on-task] × Block | 0.0023 | 0.0010 | 0.0004 – 0.0042 | 2.3597 | 6044.7236 | **0.018** |
| (Triplet Type [Low] × No-Go) × MW [on-task] | 0.0187 | 0.0045 | 0.0099 – 0.0275 | 4.1706 | 13118.9250 | **<0.001** |
| (Triplet Type [Low] × No-Go) × Block | -0.0015 | 0.0044 | -0.0100 – 0.0071 | -0.3341 | 11382.7275 | 0.738 |
| (Triplet Type [Low] × MW [on-task]) × Block | -0.0001 | 0.0008 | -0.0018 – 0.0015 | -0.1666 | 13038.3510 | 0.868 |
| (No-Go × MW [on-task] × Block | 0.0086 | 0.0047 | -0.0006 – 0.0177 | 1.8352 | 12515.6426 | 0.067 |
| (Triplet Type [Low] × No-Go × MW [on-task] × Block | -0.0015 | 0.0043 | -0.0100 – 0.0070 | -0.3518 | 13286.0048 | 0.725 |
| **Random Effects** | | | | | | |
| σ^2^ | 0.01 | | | | | |
| τ_00_ _ID_ | 0.00 | | | | | |
| τ_11_ _ID.Block_ | 0.00 | | | | | |
| τ_11_ _ID.MW [on-task]_ | 0.00 | | | | | |
| τ_11_ _ID.Triplet Type [Low]_ | 0.00 | | | | | |
| ρ_01_ | 0.61 | | | | | |
|  | 0.78 | | | | | |
|  | 0.21 | | | | | |
| ICC | 0.26 | | | | | |
| N _ID_ | 240 | | | | | |
| Observations | 14340 | | | | | |
| Marginal R^2^ / Conditional R^2^ | 0.062 / 0.303 | | | | | |

*Note.* Worse No-Go performance was associated with less accurate performance on the CTT (main effect of No-Go). The effect of No-Go was stronger during MW (interaction between No-Go and MW). Statistical learning (higher accuracy for high-probability triplets) was observed during periods of mean and good No-Go performance, with a more pronounced effect during periods of MW as No-Go performance improved (interaction between MW, No-Go, and Triplet Type).

**S3. Models on Q2 responses – Mind blanking vs. mind wandering**

***S3.1 Development of mind blanking vs. mind wandering during the task***

**Table S8.**

| *Predictor* | *Estimate* | *SE* | *t-value* | *p-value* |
| --- | --- | --- | --- | --- |
| (Intercept) | 2.28 | 0.06 | 36.81 | **<0.001** |
| block number | 0.01 | 0.00 | 1.66 | 0.096 |
| Observations | 2060 | | | |
| R^2^ / R^2^ adjusted | 0.01 / 0.001 0.02 | | | |

*Note. The relationship between the progression of the task and Q2 responses. The mean values of Q2 responses did not change as the task progressed.*

**Table S9.**

| *Predictor* | *Estimate* | *SE* | *t-value* | *p-value* |
| --- | --- | --- | --- | --- |
| (Intercept) | 0.58 | 0.02 | 25.82 | **<0.001** |
| block number | -0.00 | 0.00 | -1.30 | 0.206 |
| Observations | 30 | | | |
| R^2^ / R^2^ adjusted | 0.057 / 0.023 | | | |

*Note. The relationship between the progression of the task and the proportion of participants engaging in mind blanking. The proportion of participants engaging in mind blanking did not change as the task progressed.*

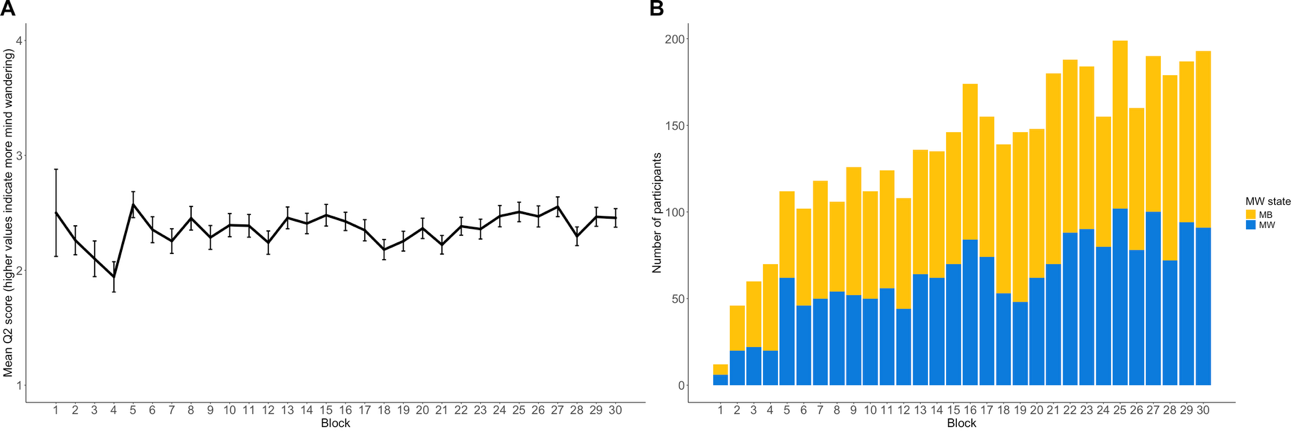

**Figure S1.** **Change in mind blanking throughout the CTT.** (A) Mean Q2 score per block, as reported by participants. The x-axis represents block number, and the y-axis shows the average Q2 score (on a scale of 1–4, where lower scores indicate more mind blanking). Error bars indicate SEM. (B) Number of participants engaged in mind blanking vs. MW per block. The x-axis indicates block number, while the y-axis reflects the number of participants. Stacked bars differentiate between participants who reported mind blanking (blue) and those who reported MW (yellow).

***S4.2 Response inhibition during periods of mind blanking vs. mind wandering***

**Table S10.**

| *Terms* | *b* | *SE b* | *95% CI* | *t* | *df* | *p* |
| --- | --- | --- | --- | --- | --- | --- |
| (Intercept) | 0.6055 | 0.0130 | 0.5800 – 0.6311 | 46.7463 | 205.9233 | **<0.001** |
| MB [MW] | -0.0086 | 0.0050 | -0.0184 – 0.0012 | -1.7401 | 125.0861 | 0.084 |
| Block | -0.0520 | 0.0059 | -0.0636 – -0.0405 | -8.8768 | 150.5885 | **<0.001** |
| MB [MW] × Block | 0.0039 | 0.0047 | -0.0054 – 0.0132 | 0.8273 | 136.5416 | 0.410 |
| **Random Effects** | | | | | | |
| σ^2^ | 0.02 | | | | | |
| τ_00_ _ID_ | 0.03 | | | | | |
| τ_11_ _ID.Block_ | 0.00 | | | | | |
| τ_11_ _ID. MB [MW]_ | 0.00 | | | | | |
| τ_11_ _ID.Block MB [MW]_ | 0.00 | | | | | |
| ρ_01_ | 0.21 | | | | | |
|  | 0.08 | | | | | |
|  | -0.10 | | | | | |
| ICC | 0.63 | | | | | |
| N _ID_ | 214 | | | | | |
| Observations | 4090 | | | | | |
| Marginal R^2^ / Conditional R^2^ | 0.043 / 0.648 | | | | | |

*Note.* No-Go performance did not differ between periods of mind blanking and MW (lack of MB or MB × Block effect).

***S3.3 Model on RT***

**Table S11.**

| *Terms* | *b* | *SE b* | *95% CI* | *t* | *df* | *p* |
| --- | --- | --- | --- | --- | --- | --- |
| (Intercept) | 19.7716 | 0.0839 | 19.6062 – 19.9370 | 235.7201 | 195.7173 | **<0.001** |
| Triplet Type [Low] | -0.1049 | 0.0151 | -0.1344 – -0.0754 | -6.9668 | 3502.6469 | **<0.001** |
| No-Go | 1.5925 | 0.0954 | 1.4054 – 1.7795 | 16.6896 | 3763.0066 | **<0.001** |
| MB [MW] | -0.0062 | 0.0197 | -0.0453 – 0.0329 | -0.3132 | 105.7217 | 0.755 |
| Block | -0.2088 | 0.0329 | -0.2738 – -0.1438 | -6.3567 | 127.5651 | **<0.001** |
| Triplet Type [Low] × No-Go | 0.3730 | 0.0889 | 0.1987 – 0.5473 | 4.1954 | 3508.3898 | **<0.001** |
| Triplet Type [Low] × MB [MW] | 0.0070 | 0.0151 | -0.0225 – 0.0365 | 0.4626 | 3503.6357 | 0.644 |
| No-Go × MB [MW] | 0.2038 | 0.0962 | 0.0152 – 0.3924 | 2.1184 | 3847.8238 | **0.034** |
| Triplet Type [Low] × block | -0.0232 | 0.0152 | -0.0530 – 0.0067 | -1.5217 | 3503.4944 | 0.128 |
| No-Go × Block | 0.0604 | 0.0948 | -0.1255 – 0.2463 | 0.6373 | 3746.0593 | 0.524 |
| MB [MW]× Block | -0.0124 | 0.0191 | -0.0499 – 0.0250 | -0.6510 | 2309.3744 | 0.515 |
| (Triplet Type [Low] × No-Go) × MB [MW] | 0.0509 | 0.0889 | -0.1233 – 0.2251 | 0.5729 | 3513.9458 | 0.567 |
| (Triplet Type [Low] × No-Go) × Block | 0.1900 | 0.0846 | 0.0242 – 0.3558 | 2.2467 | 3512.0160 | **0.025** |
| (Triplet Type [Low] MB [MW]) × Block | -0.0012 | 0.0152 | -0.0310 – 0.0287 | -0.0781 | 3502.3964 | 0.938 |
| (No-Go × MB [MW] × Block | 0.3386 | 0.0941 | 0.1542 – 0.5230 | 3.5994 | 3268.4886 | **<0.001** |
| (Triplet Type [Low] × No-Go × MB [MW] × Block | 0.0204 | 0.0846 | -0.1454 – 0.1861 | 0.2407 | 3510.0114 | 0.810 |
| **Random Effects** | | | | | | |
| σ^2^ | 0.85 | | | | | |
| τ_00_ _ID_ | 1.40 | | | | | |
| τ_11_ _ID.Block_ | 0.13 | | | | | |
| τ_11_ _ID. MB [MW]_ | 0.00 | | | | | |
| ρ_01_ | -0.04 | | | | | |
|  | -0.08 | | | | | |
| ICC | 0.64 | | | | | |
| N _ID_ | 214 | | | | | |
| Observations | 4090 | | | | | |
| Marginal R^2^ / Conditional R^2^ | 0.070 / 0.669 | | | | | |

*Note.* Participants exhibit less accurate responses during periods of MB in case of weak response inhibition (No-Go and MB interaction). As the task advances, during MB intervals in block with poor response inhibition, individuals exhibit increasing speed compared to MW blocks (interaction between No-Go, MB, and Block).

***S3.4 Model on accuracy***

**Table S12.**

| *Terms* | |  | *b* | | *SE b* | *95% CI* | *t* | *df* | *p* |
| --- | --- | --- | --- | --- | --- | --- | --- | --- | --- |
| (Intercept) | |  | 0.9163 | | 0.0053 | 0.9059 – 0.9266 | 173.9570 | 214.0299 | **<0.001** |
| Triplet Type [Low] | |  | 0.0036 | | 0.0017 | 0.0003 – 0.0070 | 2.1501 | 195.3371 | **0.033** |
| No-Go | |  | 0.1317 | | 0.0095 | 0.1130 – 0.1504 | 13.8039 | 3389.1428 | **<0.001** |
| MB [MW] | |  | -0.0017 | | 0.0023 | -0.0062 – 0.0029 | -0.7312 | 133.1783 | 0.466 |
| Block | |  | -0.0031 | | 0.0023 | -0.0077 – 0.0014 | -1.3572 | 178.4743 | 0.176 |
| Triplet Type [Low] × No-Go | |  | 0.0478 | | 0.0091 | 0.0299 – 0.0657 | 5.2341 | 3486.2581 | **<0.001** |
| Triplet Type [Low] × MB [MW] | |  | -0.0007 | | 0.0016 | -0.0037 – 0.0024 | -0.4131 | 1660.6296 | 0.680 |
| No-Go × MB [MW] | |  | 0.0232 | | 0.0097 | 0.0042 – 0.0421 | 2.3999 | 3672.5237 | **0.016** |
| Triplet Type [Low] × Block | |  | 0.0010 | | 0.0016 | -0.0021 – 0.0041 | 0.6455 | 3419.3027 | 0.519 |
| No-Go × Block | |  | 0.0296 | | 0.0095 | 0.0110 – 0.0482 | 3.1219 | 3540.6434 | **0.002** |
| MB [MW]× Block | |  | 0.0008 | | 0.0018 | -0.0028 – 0.0044 | 0.4512 | 1524.1504 | 0.652 |
| (Triplet Type [Low] × No-Go) × MB [MW] |  | | -0.0124 | | 0.0092 | -0.0304 – 0.0055 | -1.3552 | 3572.2614 | 0.175 |
| (Triplet Type [Low] × No-Go) × Block | |  | -0.0019 | | 0.0088 | -0.0191 – 0.0153 | -0.2129 | 3118.4449 | 0.831 |
| (Triplet Type [Low] × MB [MW]) × Block | |  | -0.0000 | | 0.0016 | -0.0031 – 0.0031 | -0.0125 | 3369.1103 | 0.990 |
| (No-Go × MB [MW] × Block | |  | 0.0099 | 0.0094 | | -0.0086 – 0.0285 | 1.0528 | 3312.9052 | 0.293 |
| (Triplet Type [Low] × No-Go × MB [MW] × Block | |  | -0.0081 | | 0.0087 | -0.0252 – 0.0090 | -0.9260 | 3462.7970 | 0.355 |
| **Random Effects** | |  |  |  |  |  |  |  |  |
| σ^2^ | |  | 0.01 | | | | | | |
| τ_00_ _ID_ | |  | 0.00 | | | | | | |
| τ_11_ _ID.block_ | |  | 0.00 | | | | | | |
| τ_11_ _ID.mw_2_dich1_ | |  | 0.00 | | | | | | |
| τ_11_ _ID.triplet_type1_ | |  | 0.00 | | | | | | |
| ρ_01_ | |  | 0.75 | | | | | | |
|  | |  | -0.05 | | | | | | |
|  | |  | 0.52 | | | | | | |
| ICC | |  | 0.39 | | | | | | |
| N _ID_ | |  | 214 | | | | | | |
| Observations | |  | 4090 | | | | | | |
| Marginal R^2^ / Conditional R^2^ | |  | 0.045 / 0.417 | | | | | | |

*Note.* Participants exhibit less accurate responses during periods of MB in case of weak response inhibition (interaction between No-Go and MB).

**S4. Models on Q3 responses – Spontaneous vs. deliberate mind wandering**

***S4.1 Development of spontaneous vs. deliberate MW during the task***

**Table S13.**

| *Predictor* | *Estimate* | *SE* | *t-value* | *p-value* |
| --- | --- | --- | --- | --- |
| (Intercept) | 1.80 | 0.06 | 31.46 | **<0.001** |
| block number | 0.01 | 0.00 | 3.68 | **<0.001** |
| Observations | 2060 | | | |
| R^2^ / R^2^ adjusted | 0.007 / 0.006 | | | |

*Note. The relationship between the progression of the task and Q3 responses. The overall amount of deliberate MW slightly increases as the task progresses.*

**Table S14.**

| *Predictor* | *Estimate* | *SE* | *t-value* | *p-value* |
| --- | --- | --- | --- | --- |
| (Intercept) | 0.82 | 0.02 | 39.63 | **<0.001** |
| block number | -0.01 | 0.00 | -4.40 | **<0.001** |
| Observations | 30 | | | |
| R^2^ / R^2^ adjusted | 0.409 / 0.387 | | | |

*Note. The relationship between the progression of the task and the proportion of participants engaging in deliberate/spontaneous MW. Slightly more participant engages in spontaneous MW as the task progresses.*

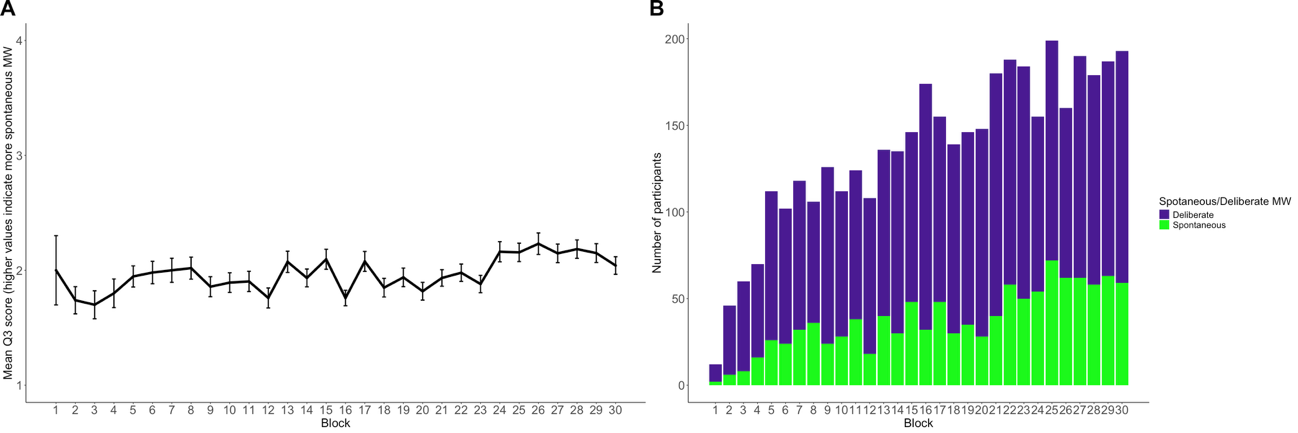

**Figure S2.** **Change in spontaneous/deliberate MW throughout the CTT.** (A) Mean Q3 score per block, as reported by participants. The x-axis represents block number, and the y-axis shows the average Q3 score (on a scale of 1–4, where lower scores indicate more spontaneous MW). Error bars indicate SEM. (B) Number of participants engaged in spontaneous vs. deliberate MW per block. The x-axis indicates block number, while the y-axis reflects the number of participants. Stacked bars differentiate between participants who reported spontaneous (green) and those who reported deliberate MW (purple).

***S4.2 Response inhibition during spontaneous vs. deliberate MW***

**Table S15.**

| *Terms* | *b* | *SE b* | *95% CI* | *t* | *df* | *p* |
| --- | --- | --- | --- | --- | --- | --- |
| (Intercept) | 0.5976 | 0.0140 | 0.5700 – 0.6251 | 42.7878 | 199.4277 | **<0.001** |
| Focus [deliberate] | 0.0157 | 0.0066 | 0.0027 – 0.0287 | 2.3846 | 119.0472 | **0.019** |
| Block | -0.0465 | 0.0063 | -0.0590 – -0.0341 | -7.4159 | 126.0548 | **<0.001** |
| Focus [deliberate] × Block | -0.0032 | 0.0054 | -0.0140 – 0.0075 | -0.5945 | 88.7134 | 0.554 |
| **Random Effects** | | | | | | |
| σ^2^ | 0.02 | | | | | |
| τ_00_ _ID_ | 0.04 | | | | | |
| τ_11_ _ID.block_ | 0.00 | | | | | |
| τ_11_ _ID.mw_3_dich1_ | 0.00 | | | | | |
| τ_11_ _ID.block:mw_3_dich1_ | 0.00 | | | | | |
| ρ_01_ | 0.14 | | | | | |
|  | -0.29 | | | | | |
|  | 0.13 | | | | | |
| ICC | 0.65 | | | | | |
| N _ID_ | 214 | | | | | |
| Observations | 4090 | | | | | |
| Marginal R^2^ / Conditional R^2^ | 0.041 / 0.661 | | | | | |

*Note.* No-Go performance is better during periods of spontaneous MW compared to deliberate MW (main effect of Focus).

***S4.3 Model on RT***

**Table S16.**

| *Terms* | *b* | *SE b* | *95% CI* | *t* | *df* | *p* |
| --- | --- | --- | --- | --- | --- | --- |
| (Intercept) | 19.7705 | 0.0855 | 19.6019 – 19.9390 | 231.3655 | 194.6841 | **<0.001** |
| Triplet Type [Low] | -0.1128 | 0.0167 | -0.1457 – -0.0800 | -6.7416 | 3547.5775 | **<0.001** |
| No-Go | 1.5321 | 0.1095 | 1.3174 – 1.7469 | 13.9895 | 2835.7576 | **<0.001** |
| Focus [deliberate] | 0.0037 | 0.0235 | -0.0429 – 0.0504 | 0.1594 | 90.0068 | 0.874 |
| Block | -0.2296 | 0.0332 | -0.2953 – -0.1639 | -6.9065 | 146.0779 | **<0.001** |
| Triplet Type [Low] × No-Go | 0.3964 | 0.1009 | 0.1985 – 0.5942 | 3.9284 | 3562.0927 | **<0.001** |
| Triplet Type [Low] × Focus [spontaneous] | 0.0172 | 0.0167 | -0.0157 – 0.0500 | 1.0255 | 3547.9310 | 0.305 |
| No-Go × Focus [deliberate] | 0.2842 | 0.1111 | 0.0664 – 0.5020 | 2.5586 | 3543.8428 | **0.011** |
| Triplet Type [Low] × Block | -0.0044 | 0.0168 | -0.0375 – 0.0286 | -0.2640 | 3547.5899 | 0.792 |
| No-Go × Block | -0.1101 | 0.1071 | -0.3201 – 0.0999 | -1.0279 | 3188.9200 | 0.304 |
| Focus [deliberate] × Block | 0.0348 | 0.0216 | -0.0076 – 0.0771 | 1.6107 | 1988.8525 | 0.107 |
| (Triplet Type [Low] × No-Go) × Focus [deliberate] | -0.0854 | 0.1009 | -0.2832 – 0.1124 | -0.8467 | 3564.2767 | 0.397 |
| (Triplet Type [Low] × No-Go) × Block | 0.2494 | 0.0955 | 0.0622 – 0.4366 | 2.6125 | 3563.0432 | **0.009** |
| (Triplet Type [Low] × Focus [deliberate]) × Block | -0.0384 | 0.0168 | -0.0714 – -0.0054 | -2.2789 | 3547.3593 | **0.023** |
| (No-Go × Focus [deliberate] × Block | 0.4399 | 0.1065 | 0.2311 – 0.6486 | 4.1317 | 2804.2346 | **<0.001** |
| (Triplet Type [Low] × No-Go × Focus [deliberate] × Block | -0.0910 | 0.0955 | -0.2781 – 0.0962 | -0.9528 | 3563.2041 | 0.341 |
| **Random Effects** | | | | | | |
| σ^2^ | 0.84 | | | | | |
| τ_00_ _ID_ | 1.43 | | | | | |
| τ_11_ _ID.Block_ | 0.12 | | | | | |
| τ_11_ _ID. Focus [deliberate]_ | 0.01 | | | | | |
| ρ_01_ | -0.02 | | | | | |
|  | -0.25 | | | | | |
| ICC | 0.64 | | | | | |
| N _ID_ | 214 | | | | | |
| Observations | 4090 | | | | | |
| Marginal R^2^ / Conditional R^2^ | 0.071 / 0.670 | | | | | |

*Note.* During spontaneous intervals, statistical learning escalates, during deliberate periods, statistical learning somewhat diminishes as the task advances (interaction between Triplet Type, Focus, and Block). During periods of spontaneous MW, RTs were faster in the case of poor No-Go performance, and slower in the case of good No-Go performance (No-Go and Focus interaction). During spontaneous periods, RTs more progressively speed up than during deliberate periods in the case of poor No-Go performance (interaction between No-Go, Focus and Block).

***S4.4 Model on accuracy***

**Table S17.**

| *Terms* | *b* | *SE b* | *95% CI* | *t* | *df* | *p* |
| --- | --- | --- | --- | --- | --- | --- |
| (Intercept) | 0.9143 | 0.0056 | 0.9033 – 0.9254 | 163.1653 | 223.7207 | **<0.001** |
| Triplet Type [Low] | 0.0046 | 0.0019 | 0.0010 – 0.0083 | 2.4941 | 242.2850 | **0.013** |
| No-Go | 0.1356 | 0.0111 | 0.1139 – 0.1573 | 12.2596 | 2987.2472 | **<0.001** |
| Focus [deliberate] | 0.0035 | 0.0027 | -0.0020 – 0.0089 | 1.2598 | 132.4354 | 0.210 |
| Block | -0.0034 | 0.0025 | -0.0083 – 0.0015 | -1.3593 | 226.3849 | 0.175 |
| Triplet Type [Low] × No-Go | 0.0499 | 0.0104 | 0.0296 – 0.0703 | 4.8075 | 3646.6612 | **<0.001** |
| Triplet Type [Low] × Focus [spontaneous] | -0.0023 | 0.0018 | -0.0057 – 0.0012 | -1.3029 | 1775.1372 | 0.193 |
| No-Go × Focus [deliberate] | -0.0097 | 0.0112 | -0.0316 – 0.0123 | -0.8635 | 3445.5846 | 0.388 |
| Triplet Type [Low] × Block | 0.0007 | 0.0017 | -0.0027 – 0.0041 | 0.3930 | 3477.2854 | 0.694 |
| No-Go × Block | 0.0155 | 0.0107 | -0.0054 – 0.0364 | 1.4571 | 2997.9907 | 0.145 |
| Focus [deliberate] × Block | -0.0006 | 0.0020 | -0.0046 – 0.0034 | -0.2888 | 1308.9905 | 0.773 |
| (Triplet Type [Low] × No-Go) × Focus [deliberate] | -0.0048 | 0.0104 | -0.0253 – 0.0156 | -0.4624 | 3516.4396 | 0.644 |
| (Triplet Type [Low] × No-Go) × Block | -0.0063 | 0.0099 | -0.0257 – 0.0130 | -0.6408 | 3347.9940 | 0.522 |
| (Triplet Type [Low] × Focus [deliberate]) × Block | 0.0003 | 0.0017 | -0.0031 – 0.0038 | 0.1970 | 3297.9302 | 0.844 |
| (No-Go × Focus [deliberate] × Block | 0.0302 | 0.0106 | 0.0093 – 0.0510 | 2.8362 | 2958.8528 | **0.005** |
| (Triplet Type [Low] × No-Go × Focus [deliberate] × Block | 0.0036 | 0.0098 | -0.0157 – 0.0229 | 0.3644 | 3615.6161 | 0.716 |
| **Random Effects** | | | | | | |
| σ^2^ | 0.01 | | | | | |
| τ_00_ _ID_ | 0.01 | | | | | |
| τ_11_ _ID.Block_ | 0.00 | | | | | |
| τ_11_ _ID.Focus [deliberate]_ | 0.00 | | | | | |
| τ_11_ _ID.Triplet Type [Low]_ | 0.00 | | | | | |
| ρ_01_ | 0.83 | | | | | |
|  | -0.35 | | | | | |
|  | 0.56 | | | | | |
| ICC | 0.39 | | | | | |
| N _ID_ | 214 | | | | | |
| Observations | 4090 | | | | | |
| Marginal R^2^ / Conditional R^2^ | 0.043 / 0.421 | | | | | |

Note. In the case of spontaneous MW, participants are getting faster only if No-Go accuracy is poor; the pattern is reversed during deliberate MW (interaction between No-Go, Focus, and Block).

**S5. Models on Q4 responses – Positive vs. negative mind wandering**

***S5.1 Development of positive vs. negative MW during the task***

**Table S18.**

| *Predictor* | *Estimate* | *SE* | *t-value* | *p-value* |
| --- | --- | --- | --- | --- |
| (Intercept) | 2.41 | 0.05 | 44.52 | **<0.001** |
| block number | 0.00 | 0.00 | 1.26 | 0.207 |
| Observations | 2060 | | | |
| R^2^ / R^2^ adjusted | 0.001 / 0.000 | | | |

*Note. The relationship between the progression of the task and Q4 responses. The mean score of the positive vs. negative dimension did not change as the task progressed.*

**Table S19.**

| *Predictor* | *Estimate* | *SE* | *t-value* | *p-value* |
| --- | --- | --- | --- | --- |
| (Intercept) | 0.52 | 0.02 | 31.93 | **<0.001** |
| block number | 0.00 | 0.00 | 0.13 | 0.902 |
| Observations | 30 | | | |
| R^2^ / R^2^ adjusted | 0.001 / -0.035 | | | |

*Note. The relationship between the progression of the task and the proportion of participants engaging in positive vs. negative MW. The proportion of participants engaging in positive vs. negative MW did not change as the task progressed.*

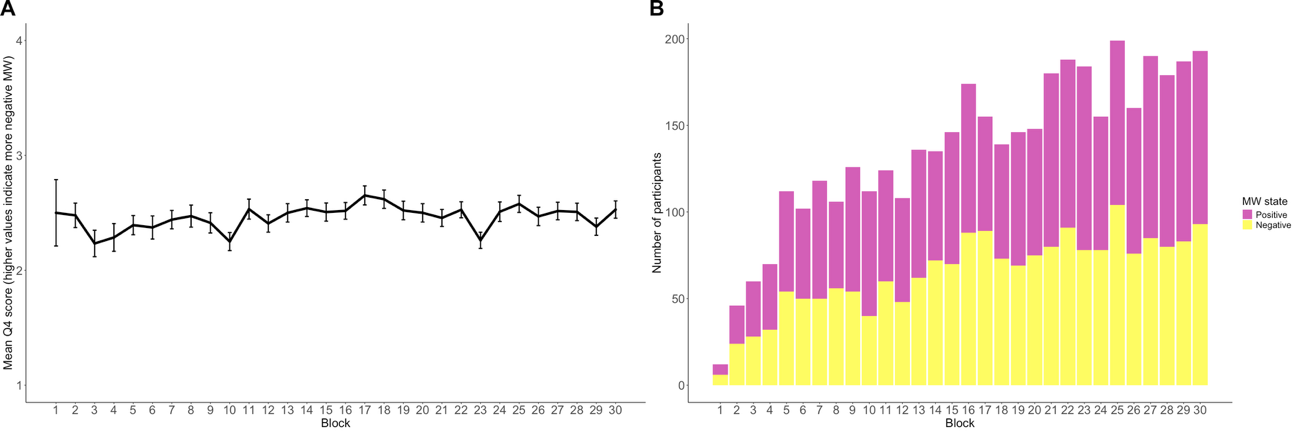

**Figure S3.** **Change in positive/negative MW throughout the CTT.** (A) Mean Q4 score per block, as reported by participants. The x-axis represents block number, and the y-axis shows the average Q4 score (on a scale of 1–4, where lower scores indicate more positive MW). Error bars indicate SEM. (B) Number of participants engaged in positive vs. negative MW per block. The x-axis indicates block number, while the y-axis reflects the number of participants. Stacked bars differentiate between participants who reported positive (pink) and those who reported negative MW (yellow).

***S5.2 Response inhibition positive vs. negative MW***

**Table S20.**

| *Terms* | *b* | *SE b* | *95% CI* | *t* | *df* | *p* |
| --- | --- | --- | --- | --- | --- | --- |
| (Intercept) | 0.6063 | 0.0131 | 0.5806 – 0.6321 | 46.4265 | 204.4427 | **<0.001** |
| Emotion [negative] | 0.0025 | 0.0054 | -0.0083 – 0.0132 | 0.4594 | 133.8610 | 0.647 |
| Block | -0.0504 | 0.0058 | -0.0618 – -0.0390 | -8.7391 | 154.4428 | **<0.001** |
| Emotion [negative] × Block | 0.0053 | 0.0047 | -0.0040 – 0.0146 | 1.1314 | 149.1638 | 0.260 |
| **Random Effects** | | | | | | |
| σ^2^ | 0.02 | | | | | |
| τ_00_ _ID_ | 0.03 | | | | | |
| τ_11_ _ID.Block_ | 0.00 | | | | | |
| τ_11_ _ID.Emotion_ | 0.00 | | | | | |
| τ_11_ _ID.Block:Emotion_ | 0.00 | | | | | |
| ρ_01_ | 0.22 | | | | | |
|  | 0.04 | | | | | |
|  | 0.26 | | | | | |
| ICC | 0.64 | | | | | |
| N _ID_ | 214 | | | | | |
| Observations | 4090 | | | | | |
| Marginal R^2^ / Conditional R^2^ | 0.040 / 0.655 | | | | | |

*Note.* No-Go performance did not differ between periods of positive and negative MW episodes (lack of Emotion or Emotion × Block effects).

***S5.3 Model on RT***

**Table S21.**

| *Terms* | *b* | *SE b* | *95% CI* | *t* | *df* | *p* |
| --- | --- | --- | --- | --- | --- | --- |
| (Intercept) | 19.7670 | 0.0839 | 19.6015 – 19.9324 | 235.6120 | 196.2313 | **<0.001** |
| Triplet Type [Low] | -0.1030 | 0.0149 | -0.1323 – -0.0737 | -6.8935 | 3603.1722 | **<0.001** |
| No-Go | 1.6129 | 0.0948 | 1.4271 – 1.7987 | 17.0228 | 3852.2465 | **<0.001** |
| Emotion [Negative] | 0.0315 | 0.0194 | -0.0064 – 0.0695 | 1.6277 | 3976.2676 | 0.104 |
| Block | -0.2137 | 0.0334 | -0.2797 – -0.1477 | -6.4035 | 128.4169 | **<0.001** |
| Triplet Type [Low] × No-Go | 0.3805 | 0.0885 | 0.2070 – 0.5540 | 4.2998 | 3606.1773 | **<0.001** |
| Triplet Type [Low] × Emotion [Negative] | -0.0214 | 0.0149 | -0.0507 – 0.0079 | -1.4308 | 3603.1233 | 0.153 |
| No-Go × Emotion [Negative] | 0.2486 | 0.0961 | 0.0602 – 0.4371 | 2.5869 | 3847.3747 | **0.010** |
| Triplet Type [Low] × Block | -0.0226 | 0.0151 | -0.0522 – 0.0069 | -1.5027 | 3603.9327 | 0.133 |
| No-Go × Block | 0.0256 | 0.0951 | -0.1609 – 0.2121 | 0.2693 | 3849.8729 | 0.788 |
| Emotion [Negative]× Block | 0.0159 | 0.0190 | -0.0213 – 0.0531 | 0.8402 | 3273.2436 | 0.401 |
| (Triplet Type [Low] × No-Go) × Emotion [Negative] | 0.0411 | 0.0884 | -0.1323 – 0.2145 | 0.4646 | 3605.5720 | 0.642 |
| (Triplet Type [Low] × No-Go) × Block | 0.2023 | 0.0853 | 0.0351 – 0.3696 | 2.3717 | 3610.9039 | **0.018** |
| (Triplet Type [Low] × Emotion [Negative]) × Block | -0.0092 | 0.0151 | -0.0388 – 0.0203 | -0.6112 | 3603.3374 | 0.541 |
| (No-Go × Emotion [Negative] × Block | 0.5055 | 0.0940 | 0.3212 – 0.6898 | 5.3774 | 3872.9210 | **<0.001** |
| (Triplet Type [Low] × No-Go × Emotion [Negative] × Block | -0.0171 | 0.0853 | -0.1843 – 0.1501 | -0.2004 | 3608.8075 | 0.841 |
| **Random Effects** | | | | | | |
| σ^2^ | 0.84 | | | | | |
| τ_00_ _ID_ | 1.40 | | | | | |
| τ_11_ _ID.Block_ | 0.14 | | | | | |
| ρ_01_ _ID_ | -0.03 | | | | | |
| ICC | 0.65 | | | | | |
| N _ID_ | 214 | | | | | |
| Observations | 4090 | | | | | |
| Marginal R^2^ / Conditional R^2^ | 0.072 / 0.671 | | | | | |

*Note.* Participants exhibit reduced speed during positive MW when they possess effective response inhibition (interaction between No-Go and Emotion).

***S5.4 Model on accuracy***

**Table S22.**

| *Terms* | *b* | *SE b* | *95% CI* | *t* | *df* | *p* |
| --- | --- | --- | --- | --- | --- | --- |
| (Intercept) | 0.9156 | 0.0053 | 0.9051 – 0.9261 | 172.4565 | 217.4791 | **<0.001** |
| Triplet Type [Low] | 0.0031 | 0.0016 | 0.0001 – 0.0062 | 2.0197 | 3626.7129 | **0.043** |
| No-Go | 0.1347 | 0.0095 | 0.1160 – 0.1534 | 14.1210 | 3354.6236 | **<0.001** |
| Emotion [Negative] | 0.0023 | 0.0021 | -0.0018 – 0.0064 | 1.1096 | 160.2574 | 0.269 |
| Block | -0.0037 | 0.0023 | -0.0082 – 0.0009 | -1.5942 | 175.2193 | 0.113 |
| Triplet Type [Low] × No-Go | 0.0483 | 0.0092 | 0.0303 – 0.0664 | 5.2528 | 3632.1408 | **<0.001** |
| Triplet Type [Low] × Emotion [Negative] | 0.0002 | 0.0016 | -0.0029 – 0.0032 | 0.1259 | 3627.7403 | 0.900 |
| No-Go × Emotion [Negative] | 0.0327 | 0.0097 | 0.0136 – 0.0518 | 3.3519 | 3850.0662 | **0.001** |
| Triplet Type [Low] × Block | 0.0010 | 0.0016 | -0.0020 – 0.0041 | 0.6637 | 3626.5120 | 0.507 |
| No-Go × Block | 0.0291 | 0.0096 | 0.0103 – 0.0478 | 3.0358 | 3595.2481 | **0.002** |
| Emotion [Negative]× Block | 0.0032 | 0.0018 | -0.0003 – 0.0067 | 1.7681 | 1597.4294 | 0.077 |
| (Triplet Type [Low] × No-Go) × Emotion [Negative] | -0.0077 | 0.0092 | -0.0257 – 0.0103 | -0.8369 | 3636.7943 | 0.403 |
| (Triplet Type [Low] × No-Go) × Block | -0.0018 | 0.0089 | -0.0191 – 0.0156 | -0.1975 | 3631.5861 | 0.843 |
| (Triplet Type [Low] × Emotion [Negative]) × Block | -0.0026 | 0.0016 | -0.0056 – 0.0005 | -1.6354 | 3626.3154 | 0.102 |
| (No-Go × Emotion [Negative] × Block | 0.0111 | 0.0095 | -0.0075 – 0.0297 | 1.1715 | 3255.3164 | 0.241 |
| (Triplet Type [Low] × No-Go × Emotion [Negative] × Block | 0.0105 | 0.0089 | -0.0069 – 0.0278 | 1.1793 | 3631.1056 | 0.238 |
| **Random Effects** | | | | | | |
| σ^2^ | 0.01 | | | | | |
| τ_00_ _ID_ | 0.01 | | | | | |
| τ_11_ _ID.Block_ | 0.00 | | | | | |
| τ_11_ _ID.Emotion [Negative]_ | 0.00 | | | | | |
| ρ_01_ | 0.80 | | | | | |
|  | -0.14 | | | | | |
| ICC | 0.38 | | | | | |
| N _ID_ | 214 | | | | | |
| Observations | 4090 | | | | | |
| Marginal R^2^ / Conditional R^2^ | 0.048 / 0.410 | | | | | |

*Note.* When No-Go performance is suboptimal, participants exhibit slightly more accuracy during negative MW. When No-Go performance is optimal, individuals exhibit increased accuracy during positive MW (interaction between No-Go and Emotion).
